# Haplotype-resolved genomes of diverse oat crown rust isolates reveal both global dispersion of long-lived clonal haplotypes and limited recombination between haplotypes

**DOI:** 10.1101/2025.08.26.672502

**Authors:** Eva C. Henningsen, Eric S. Nazareno, Rebecca Spanner, David Lewis, Jibril Lubega, Erin P. Moreau, Matthew J. Moscou, Brendan Boesen, Eric Stone, Kostya Kanyuka, Shahryar F. Kianian, Brian J. Steffenson, Peter N. Dodds, Jana Sperschneider, Melania Figueroa

**Affiliations:** Commonwealth Scientific and Industrial Research Organisation, ACT, Australia; University of Minnesota, Saint Paul, MN, U.S.A; National Institute of Agricultural Botany, Cambridge, U.K; Cereal Disease Laboratory, USDA-ARS, St. Paul, MN, U.S.A; Australian National University, ACT, Australia

**Keywords:** oat crown rust, haplotype, effectors, variation, orthology

## Abstract

Genetic diversity of pathogen populations plays an important role in host adaptation. While single high quality genome references are a valuable resource, a compilation of genome references from individuals increases the breadth of genetic variation within a species that is captured. *Puccinia coronata* f. sp. *avenae* (*Pca*), a fungal pathogen causing crown rust of oat, demonstrates rapid virulence evolution and adaptation to newly released cultivars. To broaden the geographic and temporal distribution of available *Pca* genomes, we generated nuclear haplotype-resolved genome references for ten isolates from Europe, Africa and the Middle East and compared these with existing references for USA and Australian isolates. Of the full collection of 52 haplotypes, 40 were unique. Importantly, the presence of a nearly identical haplotype in a UK isolate collected in 1984 and in USA isolates from 1990 and 2017 supports the existence of long-lived clonal haplotypes in the global population that have been exchanged between lineages. Taken together with the identification of infrequent recombination between haplotypes from geographically dispersed isolates, this evidence reflects a globally mobile population of *Pca* that is mostly comprised of persistent clonal lineages with some influence from rare recombination events. Analysis of the core and non-core proteome suggests that while the core proteome is enriched for predicted secreted and effector proteins, sequence and expression variation are most prevalent in non-core orthogroups. We anticipate that this expanded collection of haplotypes will facilitate the development of new surveillance technologies and identification of virulence loci.

## Introduction

Rust fungi are among the most destructive plant pathogens in agricultural ecosystems (Figueroa et al. 2023). As obligate biotrophs, these pathogens can only grow and reproduce in living host tissue (Lorrain et al. 2019). The life cycle of these organisms can encompass both asexual and sexual reproductive phases, which have different implications for population diversity. Some rust species require different and unrelated hosts for the sexual and asexual phases. For example, the oat crown rust fungus, *Puccinia coronata* f. sp. *avenae* (*Pca*), infects cultivated oat (*Avena sativa*) as well as related wild host plants (i.e., *Avena*, *Lolium*, *Poa* spp) during the asexual phase of its life cycle. For the sexual phase, *Pca* infects an alternate host, commonly known as buckthorn (*Rhamnus* spp.) (Simons 1970; Nazareno et al. 2018). In the absence of the alternate (sexual) host, asexual reproduction on the cereal host can occur indefinitely leading to long-lived clonal lineages. Both asexual and sexual processes contribute to host adaptation of rust fungi (Figueroa et al. 2020). Sexual recombination can introduce many new combinations of genotypes resulting in high levels of population genetic diversity and rapid emergence and spread of virulence phenotypes, such as in *Pca* populations in North America where the sexual host common buckthorn is prevalent (Nazareno et al. 2018; Miller et al. 2020). In asexual populations, the emergence of new virulence is often limited to mutation events in clonal lineages. However, the asexual stages of rust fungi are dikaryotic, meaning that each cell has two haploid nuclei and somatic exchanges of intact nuclei between two strains can also lead to new genotype combinations in the absence of the sexual stage (Figueroa et al. 2020; Duplessis et al. 2021). Such nuclear exchange events have been demonstrated to occur in multiple rust species, including wheat stem rust (*Puccinia graminis* f. sp. *tritici* - *Pgt*), wheat leaf rust (*P. triticina* - *Pt*), and *Pca* (Li et al. 2019; Sperschneider et al. 2023; Henningsen et al. 2024), and have contributed to the emergence of important epidemic strains such as the Ug99 lineage of wheat stem rust.

Virulence in rust fungi follows the gene-for-gene model first described in the flax rust pathosystem, in which the outcome of infection is determined by the recognition between resistance (R) proteins in the plant and avirulence (Avr) proteins secreted by the pathogen (Flor 1971; Dodds 2023). In the absence of recognition, these secreted proteins known as effectors facilitate infection by suppressing plant immunity and altering transcription and hormone production (Lorrain et al. 2019). A number of *Avr* genes have been identified in several rust fungi including flax rust, and cereal rust species (*Puccinia* spp.) (Chen et al. 2017; Deng et al. 2022; Lubega et al. 2024; Dodds et al. 2004; Catanzariti et al. 2005; Salcedo et al. 2017; Arndell et al. 2024). All of these encode secreted proteins that are expressed during plant infection. The emergence of virulence to specific host *R* genes in rust fungi has resulted from both *Avr* gene disruption by insertion or deletion events (e.g., *AvrSr27*, *AvrSr35* and *AvrSr50* - *Pgt*) as well as the accumulation of allelic sequence variation (e.g., *AvrL567*, *AvrP* - flax rust; *AvrRppC* - southern corn rust; *AvrSr50* - *Pgt*) (Henningsen et al. 2025). *Avr* gene expression level polymorphisms (e.g., *AvrSr27* - *Pgt*) have also been shown to have an effect on virulence (Upadhyaya et al. 2021).

The genomes of rust fungi are predicted to encode up to a few thousand effectors, with approximately 2,000 effectors annotated in Pca203 (Henningsen et al. 2022). However, the reference genome for one individual is unlikely to represent the full gene complement for the entire species. Pangenome approaches have been used in other filamentous plant pathogen species to assign gene content to core and non-core space (Badet et al. 2020; Fletcher et al. 2022; Meng et al. 2024). Non-core genes are dispensable, and in several species the non-core gene space is enriched for predicted effectors or other gene classes related to host adaptation (Fagundes et al. 2025). For example, in the fungal pathogen *Zymoseptoria tritici*, genes encoding CAZymes mostly belong to core gene space, while predicted effector genes were mostly non-core (Badet et al. 2020). However, nuclear phased genome references are only available for a handful of rust species and in most cases these are derived from only a few selected isolates and represent a limited sample of genetic diversity in the respective species. For instance, phased references are available for only four strains of *Pt*, two of *Pgt*, two of *P. striiformis* f. sp. *tritici* (*Pst* – wheat stripe rust), and just one for *P. polysora* (southern corn rust) (Wang et al. 2024; Tam et al. 2025; Liang et al. 2023; Duan et al. 2022; Sperschneider et al. 2023; Li et al. 2019). The most extensive genome reference catalogue currently is for *Pca*, which includes 16 phased, chromosome-scale reference genomes, but only representing isolates from the USA and Australia (Henningsen et al. 2022, 2024). Phylogenetic analysis of genome wide sequencing data from over 300 *Pca* isolates, representing USA, Australia, South Africa, and Taiwan, revealed high genetic diversity across populations (Miller et al. 2020; Hewitt et al. 2024; Henningsen et al. 2024). There is therefore a need to construct a more representative pangenome for *Pca* by sampling isolates from other regions and to explore sequence and expression variation to better understand host adaptation (Badet and Croll 2020).

Here we have generated nuclear phased, chromosome-scale genome references for ten *Pca* isolates sourced from Europe, Africa and the Middle East to capture additional diversity of this pathogen. The 20 haplotypes from these ten isolates bring the total *Pca* haplotype collection to 52 haplotypes, 40 of which were found to be unique. Adding to the existing evidence of movement of *Pca* haplotypes between Australia, the USA, and East Asia (Henningsen et al. 2024) an additional nuclear exchange event was identified linking *Pca* isolates from the USA and UK. Core and non-core components were established for the proteomes from unique haplotypes and allele-specific expression of single-copy heterozygous loci was explored in one isolate. The expanded haplotype-specific, globally representative atlas for *Pca* has high genotypic diversity which can be leveraged for Avr effector discovery via high throughput methods (Arndell et al. 2024).

## Results

### Expansion of the *Pca* haplotype collection and characterization of virulence phenotypes

Ten *Pca* isolates spanning sampling years from 1984 to 2018 (**Supplementary Table 1**) were chosen for phenotyping and construction of haplotype phased genome references using optimized assembly approaches (Sperschneider et al. 2023). These include seven isolates selected from an archival collection housed at the USDA-ARS Cereal Disease Laboratory and sourced from Egypt (99EGY044, 99EGY048, 99EGY050), Israel (94ISR005, 94ISR022, 94ISR045) and Switzerland (00SWI125). Two further isolates collected in South Africa (Pca43-1, Pca77-46) were selected to represent the only two genetic lineages of *Pca* detected in this country so far (Hewitt et al. 2024), while the last isolate was collected in the United Kingdom (CRO471). When combined with the published references of 16 *Pca* isolates (Henningsen et al. 2022, 2024), this collection represents isolates from seven countries across five continents (**Fig. 1A**). The isolate names and nuclear haplotype designations for the genomic references are shown in **Fig. 1B**.

**Fig. 1.**
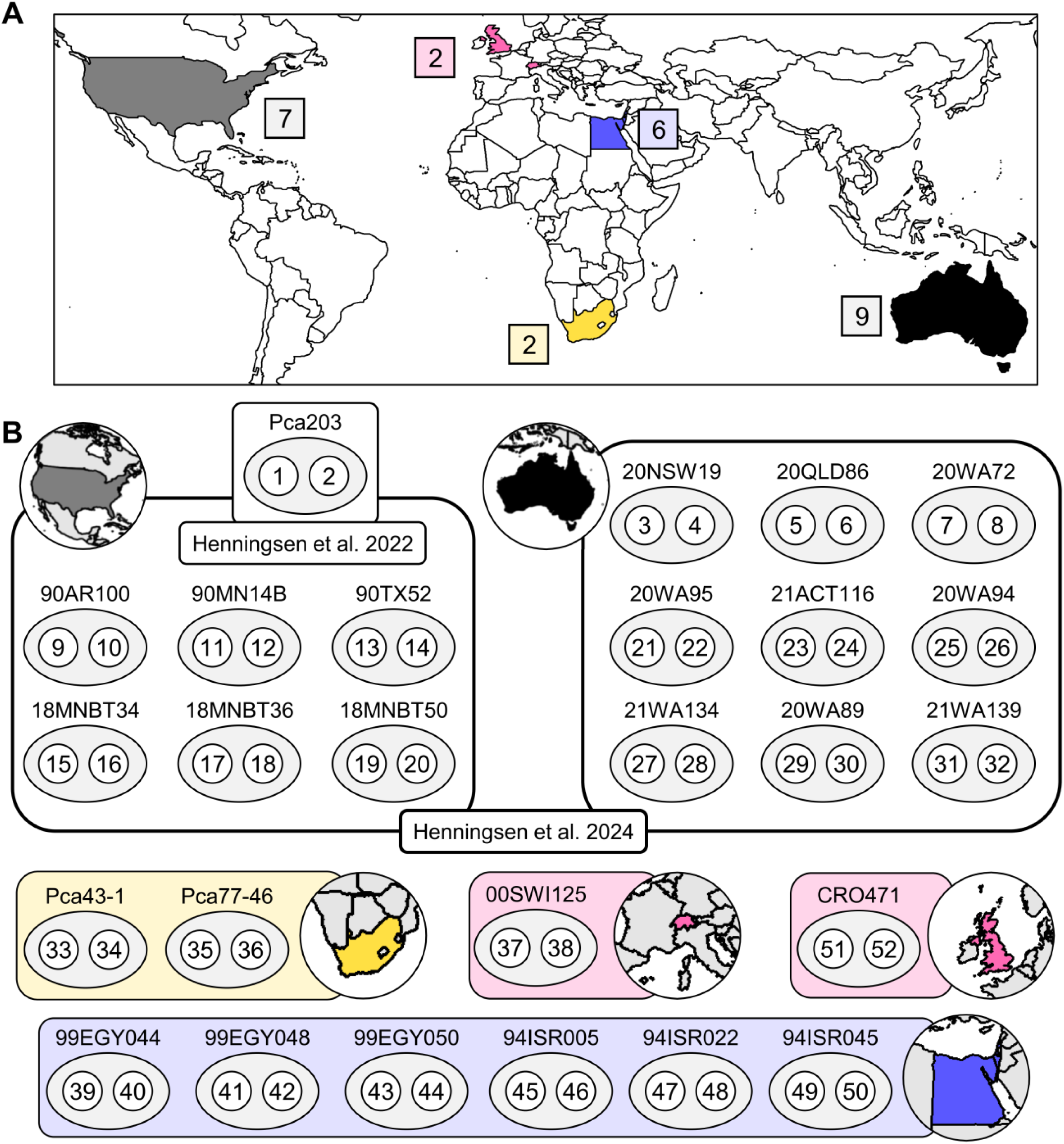
Geographic distribution of sequenced isolates and haplotype nomenclature. **A)** Map indicating sampled countries for *Puccinia coronata* f. sp. *avenae* (*Pca*) isolates with phased, chromosome level genome assemblies (published = black, grey; new = pink, yellow, blue). **B)** Isolate names and haplotype designations for published (upper two panels for USA and AUS isolates, Henningsen et al. 2024) and new *Pca* genome references (26 isolates, 52 haplotypes in total).

Virulence profiles of nine of the ten *Pca* isolates sequenced here as well as the nine Australian isolates sequenced by Henningsen et al (2024) were determined by infection of a set of 31 oat differential resistance lines. These were compared to previously determined phenotypes for the USA reference isolates (Henningsen et al. 2022; Hewitt et al. 2024), with 24 unique virulence phenotype combinations observed for these 25 isolates (**Supplementary** Fig. 1). Isolate CRO471 was phenotyped on a separate differential set with 17 common lines (**Supplementary Table 2**). One pair of isolates had identical phenotypes (99EGY044, 99EGY048), suggesting they may be closely related. The most broadly virulent isolates were from the USA (18MNBT34, 18MNBT36) and the least virulent were from Australia (21ACT116) and the USA (Pca203) (**Supplementary Table 2**). Most isolates were virulent on lines Pc45 (85%), Pc54 (74%), Pc40 (63%), and Pc46 (63%) (**Supplementary** Fig. 1**; Supplementary Table 2**). None of the isolates were virulent on Pc94, and virulence to Pc50 was only found in two isolates (20QLD86, 18MNBT50).

PacBio HiFi and Hi-C sequencing data was generated for the ten new *Pca* isolates and used to assemble nuclear-phased genomes with hifiasm (**Supplementary Table 1**). Only 11 phase swaps between four pairs of haplotypes required correction after hifiasm assembly, and the final assemblies showed 79.7-98.7% of Hi-C *trans* contacts occurring between chromosomes within the same haplotype (**Supplementary Table 3**), indicating that nuclear phasing was effective. Unplaced contigs in each assembly were mostly small (L50: 40.26-88.68 kb) and highly repetitive (67.00-97.67%; **Supplementary Table 4**). BUSCO completeness was consistent across haplotypes (95.0-95.7%) with minimal duplication (1.1-1.6%) (**Supplementary Table 4**). Continuing the established haplotype nomenclature (Henningsen et al. 2024), the new nuclear haplotypes were numbered 33 to 52 (**Fig. 1B**).

For each haplotype, scaffolding of contigs resulted in 18 chromosomes of similar sizes (3.3-8.0 Mbp) to those in previous genome assemblies (**Supplementary** Fig. 2). Chromosome 9 showed a bimodal size distribution due to the presence of a large allele-specific repetitive region adjacent to the bi-allelic *a* mating-type locus on this chromosome (Henningsen et al. 2024). Overall, synteny of chromosomes is largely preserved across all haplotypes (**Supplementary** Fig. 3), although a number of structural variants (inversions, duplications or translocations) were detected in some haplotypes, the largest of which being a 715 Kbp inversion on chr1 in hap10 (90AR100). Repetitive sequences and genes were annotated on the 20 new haplotypes which showed similar repeat (56.62-59.63%) and gene content (18,749-21,151 genes) with between 1,412 and 1,504 predicted secreted effectors per haplotype (**Supplementary Table 3**).

### Identification of an atypical extra chromosome in *Pca* haplotype 46

An additional long scaffold of 2.17 Mbp in length (referred to as Scaff1) was present in the final assembly of hap46 (94ISR005), which did not show synteny with any regions of the 18 main chromosomes of any of the *Pca* isolate haplotypes (**Supplementary** Fig. 4). Scaff1 was assembled by hifiasm as a single contig (original ID: h1tg000033l) with telomere repeat sequences at both ends and shows a centromeric contact signal (**Supplementary** Fig. 5A**, 5B**), suggesting it may represent a supernumerary chromosome. To further investigate this, an additional assembly for 94ISR005 was completed with HiCanu, using only HiFi reads. This resulted again in an identical contig with telomeres on both ends. HiFi read depth coverage of Scaff1 (50X read depth across 100.0% of Scaff1) was similar to that of the main chromosomes in 94ISR005 (47-58X) (**Supplementary** Fig. 6), while chromatin contact information clearly assigned it to hap46 and not hap45 (**Supplementary** Fig. 5A,C), together confirming that it represents nuclear genome content in one haplotype of this isolate.

Overall repeat coverage of Scaff1 (59.37%) is similar to that of other chromosomes (average =58.16%, range = 54.10 to 64.38%). Using a repeat library derived from only the main chromosomes of 94ISR005 to mask repeats on Scaff1 resulted in 58.42% bases masked, indicating that the repeat sequences in Scaff1 are shared with the rest of the *Pca* genome. Repeat family representation in Scaff1 is similar to the main chromosomes of hap45 and hap46 (**Supplementary Table 5, Supplementary** Fig. 7). Repeats on Scaff1 are slightly enriched for more diverged (older) elements compared to other chromosomes, based on Kimura distance estimation of LTRs (**Supplementary** Fig. 8). A total of 556 genes were annotated on Scaff1, including seven encoding predicted effectors, but no BUSCOs were identified.

### *Pca* haplotype comparison identifies nuclear exchanges and limited recombination

To assess the relationships among the entire set of *Pca* haplotypes a Maximum Likelihood (ML) phylogenetic tree was generated from 312,889 SNPs using hap1 (Pca203) as the reference (**Fig. 2A**). This analysis identified clonal relationships between some sets of isolates or haplotypes. For instance, the three isolates from Egypt represent a distinct clonal lineage, as the haplotypes from each isolate are near identical to each other (3,675-9,575 SNPs; hap39=hap42=hap44, hap40=hap41=hap43). The three isolates from Israel also represent a single clonal lineage with near identical haplotypes (5,023-8,046 SNPs; hap45=hap47=hap50, hap46=hap48=hap49). Interestingly, the additional 2.17 Mbp chromosome (Scaff1) in hap46 of 94ISR005 is not present in the assemblies of 94ISR022 and 94ISR045 despite the high similarity in the rest of their genomes. Mapping HiFi reads from these isolates to the 94ISR005 assembly showed only 0.0-0.1X read coverage across 0.1-2.0% of Scaff 1 (**Supplementary** Fig. 6**; Supplementary Table 6**), indicating that this additional chromosome is unique to 94ISR005 and suggesting recent acquisition or loss of this element within this clonal lineage.

**Fig. 2.**
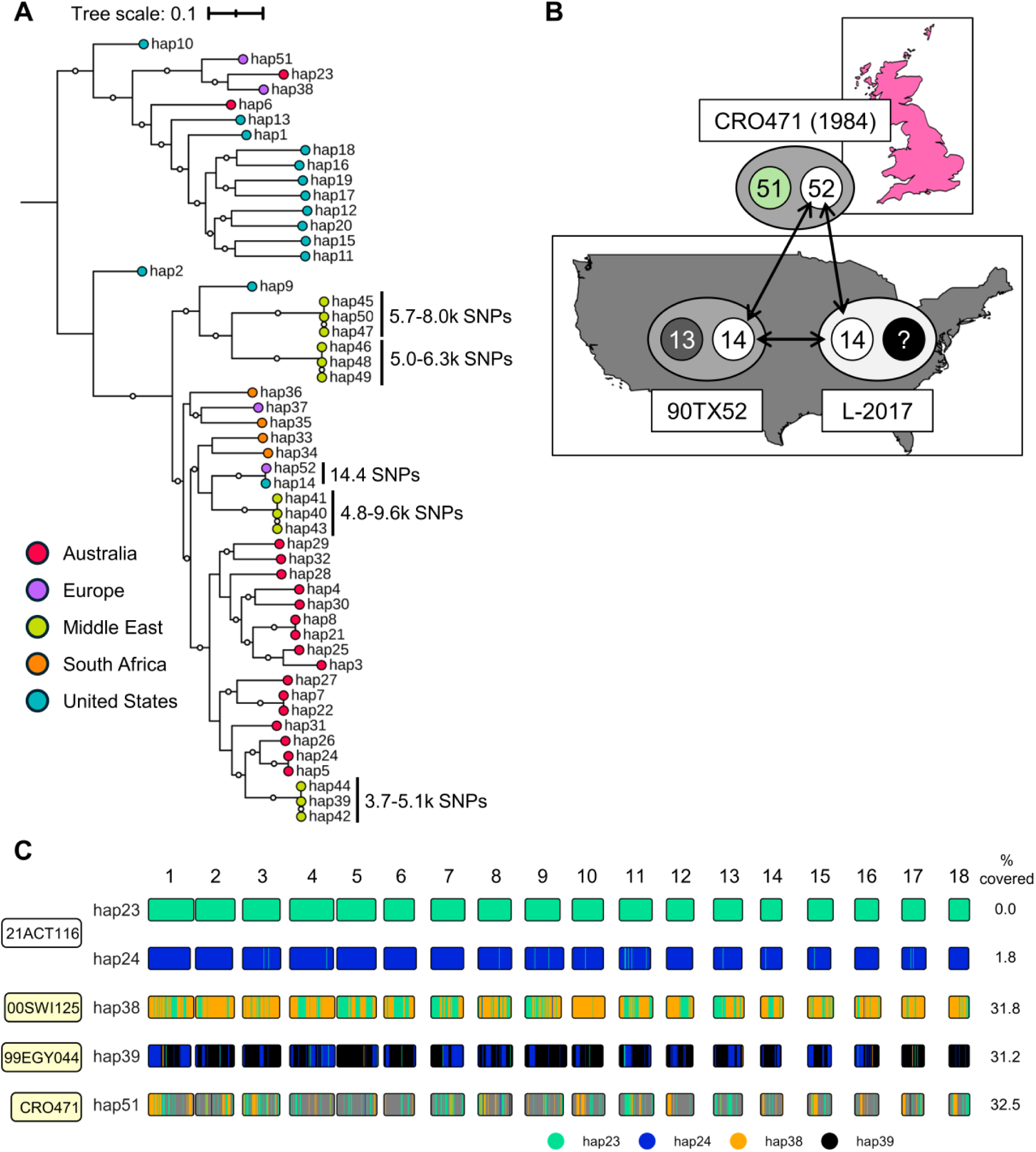
Relationships between *Pca* haplotypes. **A)** Maximum Likelihood phylogenetic tree of 52 *Pca* haplotypes constructed from 312,889 SNPs called against hap1 from Pca203. Circle fill color at branch ends reflect geographic origin of each haplotype as indicated in the key. White circles at branch midpoints represent bootstraps ≥ 80% (500 cycles). Tree scale represents mean substitutions per site. SNPs between hap14 and hap52 are reported from the vcf where hap14 was the reference. **B)** Diagram of the relationship between haplotypes from 90TX52 (hap13, hap14), CRO471 (hap51, hap52) and the previously postulated hybrid lineage L-2017 (Henningsen et al. 2024). **C)** Recombination blocks between five *Pca* haplotypes (hap23, hap24, hap38, hap39, hap51). 100 kb bins with <= 50 SNPs were considered shared between haplotypes and were assigned hierarchically from top to bottom. Chromosome fill color represents unassigned regions or those shared with haplotypes earlier in the hierarchy.

Another pair of near identical haplotypes (14,417 SNPs) were found in 90TX52-1 (hap14) and CRO471 (hap52) (**Fig. 2A; Supplementary Table 7**), while the other nuclear haplotypes in these isolates (hap13 and hap51, respectively) are highly dissimilar (372,341 SNPs). This indicates that these isolates share a common haplotype due to a nuclear exchange event and that there has been migration of a lineage containing hap14 between the US and UK (**Fig. 2B**). Notably, hap14 was previously postulated to be involved in somatic hybridization within the USA *Pca* population as it was fully contained in isolates from lineage L-2017 (Henningsen et al. 2024) (**Fig. 2B**). As noted previously (Henningsen et al. 2024), hap5 and hap24 in 20QLD86 and 21ACT116 respectively are also near identical (12,473 SNPs) and represent another nuclear exchange event within the Australian *Pca* population.

We also screened the new unique *Pca* haplotypes for blocks of shared sequence segments within chromosomes indicative of genetic recombination (Henningsen et al. 2024). Most of the new unique haplotypes (hap33-hap37, 45/47/50, hap46/48/49, hap40/41/43) do not show evidence of recent recombination with any other haplotypes captured in the collection (**Supplementary Table 8**). The only notable signatures of recombination occur within two groups of haplotypes (group 1 including hap23, hap38, and hap51, and group 2 including hap24/5 and hap39/42/44). Hap38 from 00SWI125 is 31% covered by haplotype blocks shared with hap23 from Australian isolate 21ACT116, while hap51 (CRO471, UK) is covered 32% by nonoverlapping regions of hap23 and hap38 (**Fig. 2C**). This suggests that these isolates are related by sexual recombination events and are likely separated by only 2-3 sexual cycle generations. Similarly, hap39 from 99EGY044 has 30% of its sequence covered by shared haplotype blocks with hap24 from Australian isolate 21ACT116. Hap14 (≈hap52), shared by 90TX52 and CRO471, was previously found to be mostly unrelated to other USA haplotypes, while hap13 (90TX52) was covered nearly 50% by other USA haplotypes (Henningsen et al. 2024). Given that both CRO471 haplotypes are not derived from the USA population (**Supplementary** Fig. 9A), while the unique haplotype from 90TX52 is of USA origin (**Supplementary** Fig. 9A), it is likely that hap14 was introduced into the USA from elsewhere (**Supplementary** Fig. 9B). Altogether these findings support movement of *Pca* isolates between Europe, Africa, the USA, and Australia, although the timing and direction of other migration/gene flow events cannot be inferred from this evidence (**Supplementary** Fig. 9B).

Published whole genome short read Illumina data from 352 *Pca* isolates were assessed for *k*-mer containment of the 20 new haplotypes. As expected, hap33 and hap34 from Pca43-1 were fully contained by other isolates in the same clonal lineage from South Africa (**Supplementary Table 9**). Similarly, hap35 and hap36 from Pca77-46 were contained by the other two isolates (PCA16-15-1-1, Pca2016-15-1) in the same lineage (**Supplementary Table 9**). Previously characterized clonal and putative hybrid isolates containing hap14 (90TX52-1) also had containment for hap52, in agreement with hap14 and hap52 being nearly identical (**Fig. 2A,B**). None of the other new haplotypes were contained in the 352 *Pca* isolates with short-read data. *K-*mer containment analysis also indicated that Scaff1 was not present (*k*-mer identity: 96.65-97.38%; shared *k*-mers: 48.85-57.27%) in any of the 352 *Pca* isolates with published short-read data.

### Heterozygosity maintained at both mating type loci

All ten *Pca* isolates had one copy each of the *STE3.2.2* and *STE3.2.3* alleles of the *a* mating type locus on chromosome 9, one in each nuclear haplotype (**Supplementary** Fig. 10), supporting previous findings of heterozygosity at this locus (Henningsen et al. 2024). At the *b* mating type locus, eight novel alleles (designated *bW14bE14* through *bW21bE21*) and three known alleles (*bW7bE7*, *bW11bE11*, *bW12bE12*) were identified across the 20 new haplotypes (**Supplementary** Fig. 11A,B**; Supplementary Table 10**). The ten isolates were also all heterozygous at the *b* locus. The *b* locus alleles were the same in identical haplotypes (hap14/hap52, hap45/47/50, hap46/48/49, hap39/42/44, hap40/41/43).

Protein sequences of the different *bW* and *bE* alleles were aligned to define the variable domain, DNA binding motif, and conserved domain. The N-terminal ∼125 aa of *bW* and ∼160 aa of *bE* show low conservation (mean 69% and 73% identity to consensus) across alleles and likely represent the variable domains responsible for heterodimerization (**Supplementary** Fig. 11C). The remainder of the *bW* and *bE* proteins have much less variation across alleles (mean 90% and 97% identity to consensus) and contain the universally-conserved WFXNXR nucleotide binding motifs (*bW* – WFRNAR; *bE* - WFCNAR).

### Identifying core and non-core components of the *Pca* pangenome

To identify core and non-core genes in the *Pca* pangenome we conducted an orthogroup analysis on the proteome derived from the longest isoform of all gene annotations in the 40 unique *Pca* haplotypes (**Fig. 3A**). Most of these 768,173 proteins (258,705 unique sequences) clustered into 25,833 orthogroups that contained at least two members, with just 4,555 singletons (**Supplementary Table 11**). Pangenome completeness analysis (**Supplementary** Fig. 12) indicated that cluster variation is closed (Heap’s α > 1), which agrees with the findings of Henningsen et al (2024). As estimated from the Chao lower bound (Chao 1987), the total number of clusters in *Pca* was predicted to be 36,053, suggesting the current genomes have captured ∼85% of the gene content of the *Pca* pangenome.

**Fig. 3.**
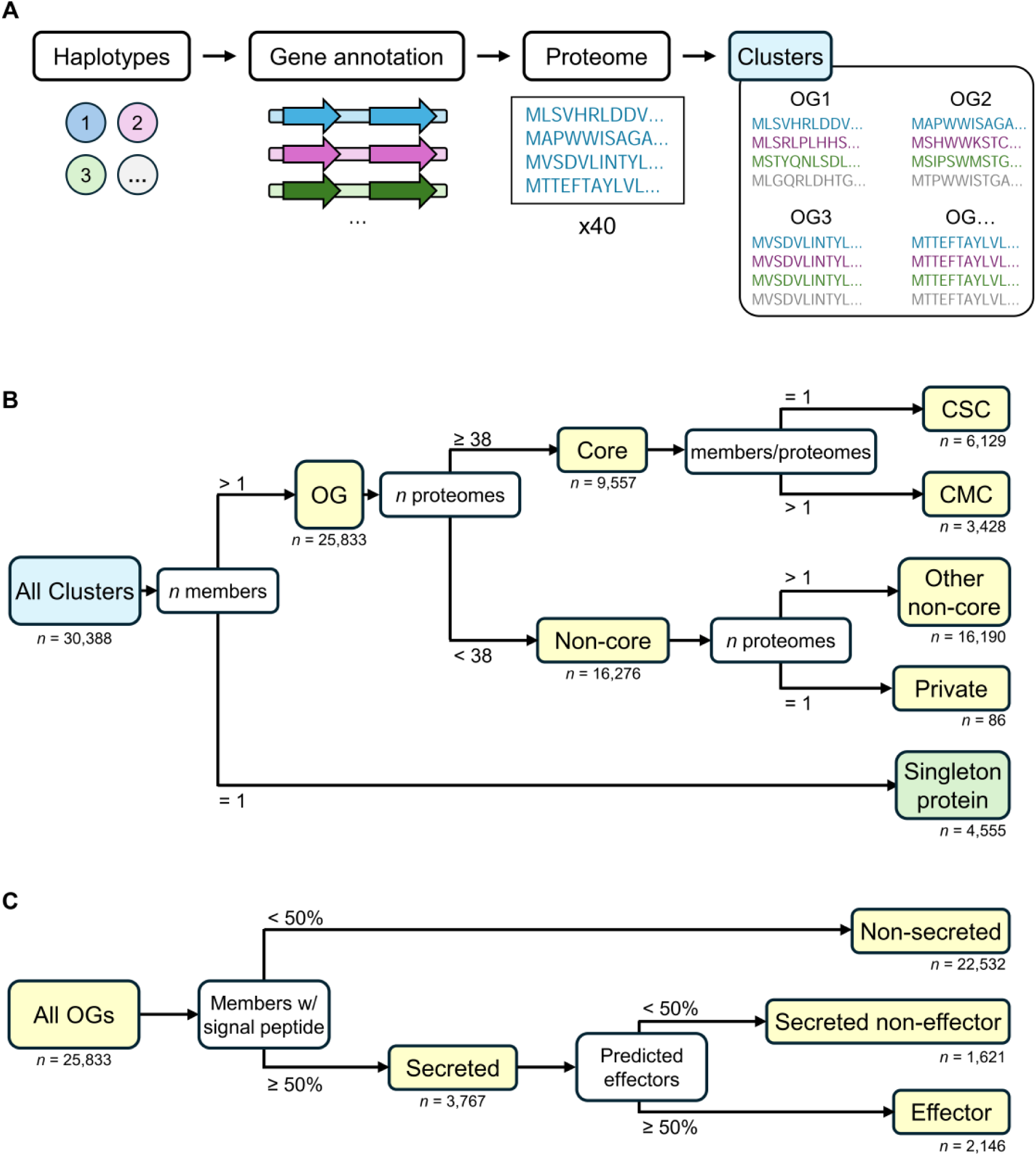
Schematic illustrating orthology analysis and orthogroup categorization. **A)** The orthology analysis begins with the set of 40 unique *Pca* haplotypes. The longest isoform of the gene annotations from all of these were translated into protein sequences, which were then clustered by sequence similarity. **B)** Clusters were divided first into orthogroups (OGs) and singleton proteins. OGs were then categorized into core and non-core groups based on their presence or absence across the proteomes. Core groups were classified as single- or multi-copy based on the copy number per haplotype and non-core groups were considered private if they occurred in a single haplotype. C) OGs were separately categorized by whether their members were predicted to be secreted or effector proteins. Yellow boxes indicate orthogroup categories, while white boxes indicate selection criteria.

The proteome of individual haplotypes consisted of between 14,083 (hap51) and 16,131 (hap10) orthogroups (mean = 15,664) with an average of 114 singleton proteins. To establish the core and non-core gene complement, the presence of orthogroup members across haplotypes was assessed (**Fig. 3B**). Orthogroups present in all 40 unique *Pca* haplotypes comprised the most frequent class (*n*=7,415; **Supplementary** Fig. 13A). However, to allow for assembly and annotation errors, orthogroups which had members from at least 38 of the 40 haplotypes (≥95%) were considered as the core component of the pangenome. This encompassed over one-third (*n*=9,557) of the orthogroups and more than half of all proteins (*n*=447,535) (**Fig. 3B**; **Supplementary** Fig. 13B; **Supplementary Table 11**). Haplotypes on average had 9,485 core (9,158 to 9,534) and 6,179 non-core (4,870 to 6,602) orthogroups. The core orthogroups were further subdivided into single copy (CSC, *n*=6,129), in which all haplotypes had exactly one ortholog, and multiple copy (CMC, *n*=3,428), in which one or more haplotypes had multiple homologs (**Fig. 3B**; **Supplementary** Fig. 13B; **Supplementary Table 11**). CMC orthogroups had between 39 and 1,189 members (mean = 59). The non-core orthogroups (*n*=16,276) ranged widely in size (2 to 758 members, mean = 19).

We also determined the representation of secreted proteins and effectors in the core and non-core components of the pangenome. Across the 40 proteomes, thirteen percent of proteins were predicted to be secreted, while seven percent were predicted to be effectors. Orthogroups were defined as representing secreted proteins or effector proteins if a minimum of 50% of members were predicted as secreted or as secreted effectors respectively (**Fig. 3C**). This resulted in 3,301 orthogroups representing secreted proteins, with 1,831 of these representing effectors (**Supplementary Table 11**). The distributions of orthogroups across haplotypes for non-secreted, secreted non-effector, and effector proteins were similar, with those present in all haplotypes being the most frequent class (**Supplementary** Fig. 13C). The core proteome was enriched for secreted and effector orthogroups relative to the non-core proteome (non-core orthogroups and singleton proteins) (χ^2^ = 206.82, df = 4, p < 2.2 x 10^-16^; **Supplementary** Fig. 13D). Non-core orthogroups with ≥ 38 members had more unique sequence variants within the group than core orthogroups (average 0.44 versus 0.31 variants per member).

### Allele-specific expression is more prevalent in gene pairs from non-core orthogroups

To examine gene expression patterns, RNAseq data were generated from *Pca* isolate 21ACT116 germinated spores (GS) and infected leaves at four or six days post infection (dpi) and used to conduct differential expression analysis (**Fig. 4A**). Of the 38,618 gene loci (18,579 orthogroups) annotated in 21ACT116, 24,427 genes (13,132 orthogroups) were expressed in samples from at least one time point (**Fig. 4A**; **Supplementary Table 12**). A much greater proportion of genes from core orthogroups (80%) were expressed relative to genes from non-core orthogroups (41%) (**Fig. 4B**). A large number of expressed genes showed differential expression (DE) between GS and the two infection timepoints (**Supplementary** Figs. 14A,B), with around one third upregulated in the infection samples relative to GS and around one fifth downregulated (**Supplementary Table 12**). However, expression profiles at 4dpi and 6dpi were very similar, with few genes showing DE between these time points (**Supplementary** Fig. 14C). A substantially higher proportion of genes from secreted and effector orthogroups were upregulated during infection (37.1-66.4%) than those from non-secreted orthogroups (23.0-34.5%), regardless of their status as core or non-core (**Fig. 4B**, **Supplementary** Fig. 15).

**Fig. 4.**
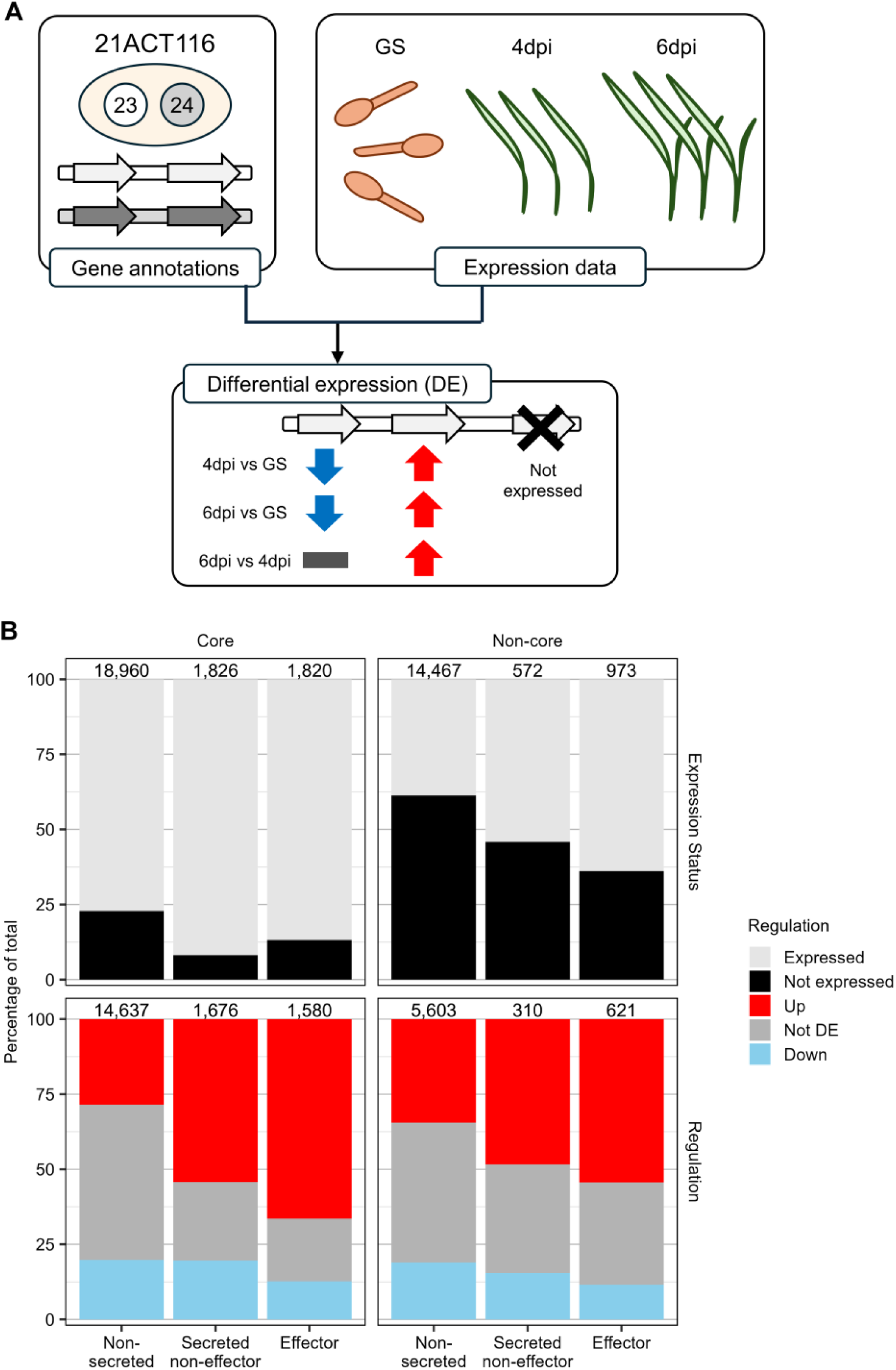
Differential expression analysis for genes from *Pca* isolate 21ACT116. A) Differential expression (DE) analysis was conducted by quantifying the RNAseq reads which map to the 38,618 annotated loci in 21ACT116 haplotypes hap23 and hap24 from three replicates for each of three timepoints (germinating spores – GS, 4 days post-inoculation – 4dpi, 6 days pos-inoculation – dpi). B) Percentage (y-axis) of genes from orthogroups in different categories (columns, x-axis) and whether they are expressed or not (top row) or differentially regulated (bottom row) for the comparison of 6dpi infected leaf samples to GS. Fill color corresponds to expression or regulation of genes: expressed (light grey), not expressed (black), downregulated (blue), not DE (grey), or upregulated (red). The total number of genes represented is shown at the top of each bar.

Virulence phenotypes can result from differences in expression between alleles of *Avr* genes, such as for *AvrSr27* in *Pgt* (Upadhyaya et al. 2021). To assess such allele-specific expression (ASE) differences in *Pca*, we first selected genes belonging to orthogroups with exactly one member from hap23 and one from hap24 (**Fig. 5A**). Of 10,897 such pairs, almost all (98.1%) consisted of genes located on homologous chromosomes representing likely allele pairs. Most of these (*n* pairs = 9,419) had sequence differences between the transcripts of the two genes, allowing for allele-specific expression analysis (**Fig. 5B**; **Supplementary Table 12**). At least one gene from most allele pairs was expressed during at least one timepoint (*n* pairs=8,157). ASE (| log_2_(fold change) | ≥ 1, p-value ≤ 0.05) was observed for 3,610 (38.3%) of the heterozygous loci in 21ACT116 within at least one timepoint, with 2,691 loci exhibiting ASE at 6dpi (**Fig. 5B**; **Supplementary Table 12**). The prevalence of ASE did not differ substantially between core or non-core orthogroups or between non-secreted, secreted non-effector, and effector orthogroups (**Fig. 5B**; **Supplementary** Fig. 16A). However, for pairs exhibiting ASE, the magnitude of DE was higher in non-core pairs than in core pairs (one-sided t-test: t = -13.427, df = 3,231.9, *p*-value = 2.44 x 10^-40^; **Fig. 5B**).

**Fig. 5.**
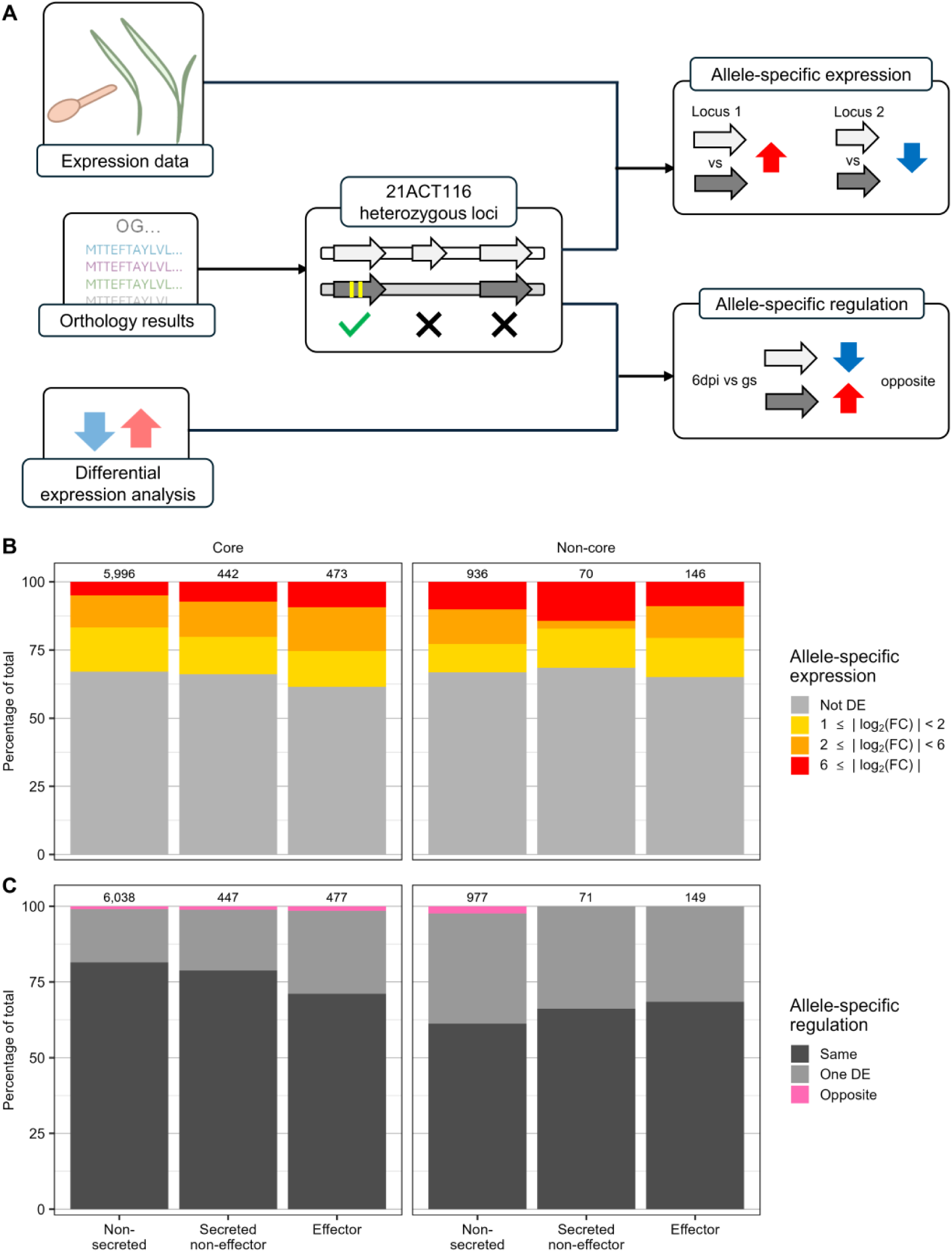
Allele-specific expression and allele-specific regulation analysis. **A)** Orthogroups were selected that contained exactly two members from 21ACT116, one each from hap23 and hap24 on homologous chromosome and with at least one sequence difference between their transcripts (*n*=9,419). To assess allele-specific expression (ASE), RNAseq data was analyzed using the orthogroup ID as the locus ID with haplotype as the treatment for DE analysis conducted separately for each of the three timepoints (germinated spores - GS, 4 days post-inoculation – 4dpi, 6 days post-inoculation – 6dpi). Allele-specific regulation (ASR) was assessed by examining the DE results from the 4dpi vs. GS and 6dpi vs. GS comparisons to pairs where the two alleles exhibited the same or different DE patterns. **B)** Percentage of expressed heterozygous loci (y-axis) from 21ACT116 in each orthogroup category (x-axis, columns) exhibiting degrees of differential expression (fill color). Categories represent ranges for the absolute value of log_2_(Fold change): Not DE - 0 to < 1), 1 to < 2; 2 to < 6); ≥ 6. **C)** Percentage of expressed allele pairs that showed ASR (y-axis) for different orthogroup categories (x-axis, columns) for the comparison of 6dpi infected leaf samples to GS. Fill color reflects what type of difference was observed (Same = both alleles had the same regulation pattern; One DE = one allele was DE, the other was not; Opposite = one allele was upregulated and the other downregulated). In panels **B** and **C**, the total number of genes represented is shown at the top of each bar.

We also examined the prevalence of allele-specific regulation (ASR), where one allele showed DE and the other did not or where one allele was upregulated and the other was downregulated (**Fig. 5A**). About a quarter of gene pairs showed ASR at 4dpi (25.5%) or 6dpi (22,6%) relative to GS, mostly as a result of one allele being DE and the other not (**Fig. 5C**, **Supplementary** Fig. 16B). ASR was observed more frequently for genes from non-core orthogroups (37.6%) than for genes from core orthogroups (19.4%) (**Fig. 5C**; **Supplementary Table 12**). Core effector genes (28.9%) also exhibited ASR more frequently than core non-effector genes (non-secreted 18.5%; secreted non-effector 21.3%) (**Fig. 5C**). ASR was slightly less prevalent in non-core effector pairs (23.5%; **Supplementary Table 12**).

## Discussion (1,241 words)

The additional ten *Pca* isolates sequenced to chromosome and haplotype level resolution have expanded the current haplotype collection to 52 haplotypes, 40 of which are unique. These represent isolates collected across six decades, seven countries and five continents. The presence of near identical haplotypes, hap14 and hap52, in a UK isolate collected in 1984 and USA isolates from 1990 and 2017 (this study and Henningsen et al. 2024) supports the existence of long-lived clonal haplotypes and nuclear exchange between lineages in the global *Pca* population. This is consistent with previous observations of somatic hybridization in the Australian *Pca* population and between Australian and Taiwan isolates (Henningsen et al. 2024) as well as the wide prevalence of nuclear exchange in other cereal rust species (Li et al. 2019; Sperschneider et al. 2023; Hovmøller et al. 2025). In addition, there is evidence for some limited recombination between globally dispersed *Pca* haplotypes, likely as a result of rare sexual reproduction. For instance, hap39 from Egyptian isolate 99EGY044 and hap24 from Australian isolate 21ACT116 share ∼30% of their sequence in large recombination blocks, suggesting that they are separated by two generations of sexual reproduction. Given these isolates were collected 22 years apart, this also suggests a long period of clonal maintenance of these haplotypes post-recombination. Similarly, hap51 from CRO471 (collected in 1984 in UK), hap38 from 00SWI125 (2000, Switzerland) and hap23 from 21ACT116 (2021, Australia) share large recombinant block sequences covering ∼30% of their genomes, again indicating a close familial relationship despite spanning 37 years in collection times. Notably, the UK isolate CRO471 contains one nucleus shared with USA isolates by nuclear exchange, and a second nucleus related by recombination to isolates collected in Europe and Australia, suggesting movement of *Pca* isolates carrying these haplotypes between Europe, Africa, USA and Australia. Isolate 21ACT116 containing hap23 and hap24 was previously postulated to represent one of the founding lineages of the Australian *Pca* population as an exotic incursion (Henningsen et al. 2024).

Overall, these observations suggest that the globally mobile population of *Pca* is largely composed of long-lived clonal lineages with some exchange of haplotypes but is also influenced by relatively rare recombination events. This contrasts with the situation within the USA, where the local *Pca* population structure is shaped by frequent sexual reproduction facilitated by the invasive alternate host *R. cathartica* and individual isolates share very small regions of common recombination blocks (Henningsen et al. 2024). Despite this, some haplotypes found in USA haplotypes are likely introduced from other *Pca* populations. For example, hap13 from 90TX52 shows strong signatures of recombination with other USA haplotypes, while hap14 of this isolate is genetically distinct from US haplotypes but is near identical to hap51 found in Europe (CRO471) and shows signatures of recombination with other international *Pca* isolates. This evidence suggests that hap14 was likely introduced to the USA by migration of an international isolate which contributed hap14 by nuclear exchange with a local strain to form the 90TX52 lineage. The *Pca* population in Australia is represented by a limited number of clonal lineages, some of which are also related by nuclear exchange events and rare recombination (Henningsen et al. 2024). For instance, hap25 in 20WA94 (representing clonal lineage 11) is an F_1_-recombinant with 50% of its sequence derived from each of hap3 and hap4 in 20NSW19 (representing clonal lineage 18).

While there are now genome references representing *Pca* isolates from Europe, Africa, and West Asia, there has not been a deep sampling of the populations from these areas for short read sequencing as was previously done for the USA, South Africa, and most recently Australia and Taiwan (Ho et al. 2024; Henningsen et al. 2024; Hewitt et al. 2024; Miller et al. 2020). In addition, there is little representation of *Pca* throughout Europe and parts of Russia where *Rhamnus* species are endemic (Simons 1985), so it is not clear to what extent *Pca* undergoes sexual reproduction in these areas. Thus, the understanding of the population-level genotypic diversity is still limited in these regions.

Eight novel *b* locus alleles were defined from the 12 new unique haplotypes, suggesting that additional allelic diversity at this locus remains undescribed in *Pca*. In contrast, only two alleles have been detected at the *a* locus, and all isolates are heterozygous at both loci.

All but one of the *Pca* isolates have 18 chromosomes, which are highly syntenous but display numerous but mostly small structural variations (inversions, deletions, duplications and translocations). The exception to this is isolate 94ISR005, which possessed an additional chromosome-like sequence in one nuclear haplotype (hap46). This element was assembled as a single 2.17 Mbp contig containing telomere sequence repeats at both ends and an internal region showed centromeric connections. Read coverage was equivalent to the rest of the genome and Hi-C data clearly placed this sequence in the hap46 nucleus, suggesting that it represents an additional chromosome present in this nucleus. This chromosome contains a similar repeat content to the rest of the genome but shows no particular homology to any other chromosome except in very short regions and is smaller in size than the main chromosomes (next smallest chromosome is ∼3.0Mbp). Rearrangements and horizontal chromosome transfer between related species are known to contribute to new chromosome formation in other filamentous plant pathogens (Langner et al. 2021; Zhang et al. 2024). Homologous chromosomes between *Puccinia* species generally maintain macrosynteny (Edwards et al. 2022; Henningsen et al. 2022), so the origin of the extra scaffold in hap46 is not clear given its lack of synteny to any of the main *Pca* chromosomes. In Ascomycete fungi, accessory chromosomes generally have high repeat content and fewer genes than core chromosomes but are enriched for effectors and are known to contribute to host adaptation (Croll and McDonald 2012; Bertazzoni et al. 2018; Barragan et al. 2024). Unlike these accessory chromosomes, the extra chromosome found in *Pca* isolate 94ISR005 has comparable repeat coverage and higher gene density but lower effector gene content relative to the main chromosomes. Interestingly, this chromosome was present in only one of three otherwise clonal isolates, suggesting recent acquisition or loss of this element within this lineage. Although isolate 94ISR005 shows a substantially different virulence profile to the other two clonal isolates, it is not clear whether this is related to the presence of the additional chromosome.

To prioritise candidate effectors for the future identification of *Avr* genes, we partitioned the *Pca* pangenome into core and non-core proteins. The core gene space conserved across the haplotypes comprises roughly half of the genes in the *Pca* pan-genome and about two thirds of the genes in each haplotype, while the remainder (non-core genes) showed presence-absence variation (PAV). Non-core genes also showed enhanced sequence variation compared to core genes. Changes in virulence can arise from changes in gene expression, as observed for *AvrSr27* in the stem rust fungus (Upadhyaya et al. 2021). We observed about 20-30% of allele pairs in isolate 21ACT116 showing either differential expression or differential regulation and this was consistent across all orthogroup classes (core and non-core, effectors and non-effectors). Altogether, our evidence suggests that changes in virulence in *Pca* could be driven by all of the mechanisms established for other rust species including PAV, copy number variation, amino acid substitutions, and gene expression changes (Chen et al. 2017; Salcedo et al. 2017; Upadhyaya et al. 2021; Ortiz et al. 2022; Chen et al. 2025). We anticipate that the most likely *Avr* gene candidates would be members of an orthogroup where most proteins are predicted as effectors with relatively high sequence variation or PAV. These criteria could be used to inform candidate selection for high-throughput *Avr* screening.

## Materials and Methods (1,764 words)

### *Pca* isolate amplification and phenotyping

All isolates except CRO471 were recovered from liquid nitrogen storage at the USDA-ARS Cereal Disease Laboratory (St. Paul, MN) and heat shocked at 45°C for fifteen minutes (**Supplementary Table 1**). Spores were then suspended in Soltrol 170 Isoparaffin oil (Chevron Phillips, TX, USA) and sprayed onto seedlings of cultivars ‘Marvellous’ or ‘Starter’ which had been previously treated with maleic hydrazide. Plants were allowed to dry for at least 40 minutes before incubation in humidity chambers overnight (99% relative humidity). After humidity treatment, plants were removed to growth chambers set to 23°C for 16 hours light and 18°C for 8 hours dark. Phenotyping was conducted at 10-11 days post inoculation (dpi) on a subset of the USA oat differential set described by Nazareno et al (2018). Susceptible infection types were recorded as “H” and resistant infection types were recorded as “L”.

Isolate CRO471 was revived from original infected leaf samples housed at National Institute of Agricultural Botany (NIAB, Cambridge, UK) by shaking infected leaves over detached leaves of susceptible oat cultivar ‘Selma’ mounted on benzimidazole agar. Spores collected from the detached leaves where then used to infect pots of ‘Selma’ grown at 22°C for 16 hours of light and at 12°C for 8 hours darkness. Spores were collected at 14 dpi. Oat lines for phenotyping were inoculated using an oil suspension of spores applied by airbrush at the second leaf stage. Phenotyping was conducted at 14 dpi.

### Sample preparation for nucleic acid sequencing

High molecular weight (HMW) DNA extracted from rust spores as described by Li et al (2019) was used to prepare PacBio HiFi libraries and sequenced to 30-50X depth at the University of Minnesota Genomics Center. Spores were prepared for Hi-C as described by Sperschneider et al. (2023). Libraries were prepared at Phase Genomics (Seattle, WA, USA) and sequenced with Illumina Novaseq by Azenta Life Sciences (South Plainfield, NJ, USA; formerly Genewiz). RNA for *Pca* isolate 20ACT40, from the same lineage (L2) as 21ACT116 (described in Henningsen et al. 2024), was obtained from germinated spores and from infected leaf tissue at 4 and 6 dpi. To obtain a germinated spore mat, freshly collected spores were dispersed over a thin layer of germination solution (0.0972% [v/v] ethanol, 0.0048% [v/v] Tween 20, 0.0008% [v/v] nonanol) and left to incubate in the dark at room temperature for 12 hours. Three replicates were collected. Infected leaf samples were obtained by pipetting 20 mg spores suspended in 400 µL Novec oil (3M, Saint Paul, MN, USA) onto first-leaf stage seedlings of oat cv. ‘Swan’. A single leaf per replicate was collected at 4dpi and 6dpi. Three separate inoculations were conducted for both timepoints as independent biological replicates. Spore mat and infected leaf samples were flash frozen immediately in liquid nitrogen and stored at -80C until RNA was extracted with the Maxwell RSC Plant RNA Kit (Promega, Madison, WI, USA) on the Maxwell RSC 48 Instrument (Promega, Madison, WI, USA). Stranded libraries were sequenced at the Australian Genomics Research Centre (AGRF) on Illumina NovaSeq to obtain 150 bp reads.

### Genome assembly pipeline

Hifiasm v0.19.5 was used to assemble PacBio HiFi reads in Hi-C integration mode (Cheng et al. 2021). The resulting haplotypes were combined and contigs were filtered for low coverage, mitochondrial gene content, and contamination as described by Sperschneider et al. (Sperschneider et al. 2023). Briefly, HiFi reads were mapped onto the combined assembly with minimap2 v2.22 (Li 2018) and contigs with less than 5X or 10X average coverage (depending on the input read coverage) were excluded. The remaining contigs were filtered for mitochondrial and contaminant contigs with blast+ v2.11.0 (Camacho et al. 2009). NuclearPhaser (https://github.com/JanaSperschneider/NuclearPhaser) was used to identify and correct any phase swaps in the assemblies as previously described (Duan et al. 2022). Scaffolding was performed by examining Hi-C contact maps created with HicPro and hicexplorer (Servant et al. 2015; Ramírez et al. 2018) and outputs from SALSA2 v2.3 (Ghurye et al. 2017). HiCanu (genomeSize=95) was used to check the assembly of 94ISR005 (Nurk et al. 2020).

### Repeat masking and annotation

Repeat sequences in full assemblies were masked by running RepeatModeler v2.0.2a (-LTRStruct) and Repeatmasker v4.1.2pl (Smit et al. 2013; Flynn et al. 2020). For statistics, all repeats were used (-gff -engine ncbi) whereas for further annotation, only classified repeats were masked and other options were selected to reduce masking of low-complexity DNA (-nolow -engine ncbi). The Kimura distance estimates for LTR repeats on the 94ISR005 chromosomes were extracted from the output of RepeatMasker (Smit et al. 2013) v.4.1.2pl run with (-gff -a) and repeat family coverage was calculated with one_code_to_find_them_all.pl (Bailly-Bechet et al. 2014).

BUSCO representation was estimated with busco3 v3.0.2 using the odb9 Basidiomycota lineage dataset (Waterhouse et al. 2018). Annotation was performed as previously described (Sperschneider et al. 2023; Henningsen et al. 2024) using published RNAseq data from *Pca* isolates 12NC29 (NCBI: PRJNA398546) and Pca203 (CSIRO DAP: 53477) (Miller et al. 2018; Henningsen et al. 2022). The Pca203 genome reference was reannotated using the described method. Secreted proteins were selected based on signal peptide predictions with SignalP v4.1 and lack of transmembrane domains predicted with TMHMM v2.0c (Möller et al. 2001; Nielsen 2017) and effectors predicted using EffectorP v3.0 (Sperschneider and Dodds 2022).

### Identification of mating type alleles

Protein sequences for *STE3.2.2*, *STE3.2.3*, *bW-HD1* and *bE-HD2* from Pca203 were BLASTed (tblastn) against the new haplotypes with blast+ v2.11.0 (Camacho et al. 2009). Protein sequences for *STE3.2*, *bW-HD1*, and *bE-HD2* alleles were aligned with CLUSTALW, and phylogenetic trees were constructed with the built-in ‘raxml-bootstrap’ option (Larkin et al. 2007). Protein trees were rooted on alleles from *Pgt* CRL 75-36-700-3 (*Pgt STE3.2.1, Pgt bW-HD1*, and *Pgt bE-HD2*) with R package ‘ape’ and visualized with iTOL (Cuomo et al. 2017; Paradis and Schliep 2019; Letunic and Bork 2021). The 5’ to 3’ DNA sequence of the *HD* loci from new haplotypes were sketched with mash v2.3 (-s 1000; Ondov et al. 2019) and published Illumina reads for *Pca* isolates were then screened against these sketches. Alleles were considered contained at or above 99.95% *k*-mer identity and 98.90% shared *k*-mers. Conservation across the *b* mating type locus protein sequences was calculated using Jalview v2.11.4.1 (Waterhouse et al. 2009). The conserved DNA binding domains (WFXNXR motif; Duboule 1994) for the *bW* and *bE* proteins were identified manually by inspecting the alignment.

### *k*-mer containment of haplotypes

*K*-mer sets for each haplotype were generated with mash sketch (-s 100000) (Ondov et al. 2019). Illumina reads for 352 *Pca* isolates were compared to the *k*-mer sets with mash screen and haplotypes were considered contained by an isolate if *k*-mer identity was >= 99.98% and shared *k*-mers were >= 99.80%. For containment analysis of Scaff1, sketch size was set to 50,000 (-s 50000).

## Haplotype comparisons

Pairwise comparisons between all published and new *Pca* haplotypes were conducted with nucmer and dnadiff from mummer v4.0.0 (Kurtz et al. 2004; Marçais et al. 2018). To assess potential recombination among haplotypes, the cactus-pangenome pipeline from cactus v2.6.6 (Hickey et al. 2024) along with custom scripts were run as previously described by Henningsen et al (2024). Non-reference SNPs from each comparison of haplotypes were counted in 100 kb interval bins and bins with less than 50 SNPs were flagged as shared. Shared haplotype blocks were assigned with an ordered hierarchy determined by calculating pairwise overlaps between all haplotypes.

For haplotype phylogenies, variants on hap1 in the other 51 *Pca* haplotypes from cactus-pangenome were filtered with vcftools v0.1.16 (Danecek et al. 2011) (--min-alleles 2 –max alleles 2 –max-missing 0.9 –maf 0.05) and a sample column for hap1 with reference calls at every site was added. The filtered vcf was converted to a phylip alignment with vcf2phylip (Ortiz 2019) and input into RAxML v8.2.12 for ML tree estimation and bootstrapping (-f a -m GTRCAT -# 500 –no-bfgs) (Stamatakis 2014). The resulting tree was visualized with iTOL (Letunic and Bork 2021).

To assess fine-scale synteny, haplotypes were aligned with mummer v4.0.0 (Marçais et al. 2018) and alignment information was processed with SyRI v1.6.3 (Goel et al. 2019) before visualization with plotsr v1.1.0 (Goel and Schneeberger 2022).

### Orthology

The longest isoform of gene annotations from the chromosomes of the 40 unique *Pca* haplotypes were used as the input protein set for Orthofinder v2.5.4 (Emms and Kelly 2019). Orthology analyses were conducted on the proteomes of haplotypes rather than isolates. Custom scripts were used to extract statistics (https://github.com/henni164/Pca_pangenome2). One-to-one orthologs from 21ACT116 were extracted by screening orthogroups where there were exactly two members from 21ACT116, one each from hap23 and hap24. The genes in each pair had to be annotated on homologous chromosomes and their transcripts had to have at least one SNP to be included in analysis.

### Pangenome saturation analysis

Pangenome saturation analysis was conducted as previously described (Henningsen et al. 2024); briefly, cluster gene counts (orthogroups and singletons) per haplotype were used as the presence/absence input for R package micropan v2.1 (Snipen and Liland 2015). micropan function ‘heaps’ was used to estimate the decay parameter α, and function ‘rarefaction’ was used to generate 1000 random permutations for the cumulative counts of orthogroups. Only the set of unique haplotypes was used (*n*=40, excluding hap5, hap21, hap22, haplotypes 41 to 44, haplotypes 47 to 50, hap52).

### Differential gene expression analysis

RNAseq reads from 20ACT40 GS and infected leaf samples (4dpi and 6dpi) were trimmed with fastp (Chen et al. 2018). Salmon quant (-l ISR) (Patro et al. 2017) was run on individual replicates of the trimmed RNAseq reads for the three timepoints after indexing (--keepDuplicates). Differential expression between sample types (6dpi versus 4dpi, 4dpi versus GS, 6dpi versus GS) was assessed with R package DESeq2 (Love et al. 2014) and visualized with R package ggplot2 (Wickham 2016). Differentially expressed genes were considered to have an adjusted p-value < 0.05 and a fold change of at least 2. Allele-specific expression between one-to-one orthologs from 21ACT116 was also assessed with DESeq2 (Love et al. 2014), using haplotype as a treatment and orthogroup ID as the locus ID.

### Map visualization

Globe-projected world map were drawn with R package ‘giscoR’ v.0.5.0 (Hernangómez 2024) which uses GISCO map data released by Eurostat. Flat world maps were constructed with the R ‘maps’ v.3.4.2 package (Becker. R.A. et al. 2023), which uses Natural Earth project data (https://www.naturalearthdata.com/) in the public domain. R package ‘ggplot2’ v3.5.1 was used in these visualizations (Wickham 2016).

## Data availability

Raw reads generated in this study are available on NCBI under BioProject accession PRJNA1289219. Genome assemblies are available on the CSIRO Data Access Portal (https://data.csiro.au/collection/csiro:65829). Code used to conduct analyses and generate figures is available on Github (https://github.com/henni164/Pca_pangenome2).

## Acknowledgements

We thank CSIRO High Performance Computing Services for computational resources, and Peter Tyson and Joel Hansen for providing computational support. We also thank Emmery Hartwig and Jakob Riddle for their assistance in amplifying the crown rust isolates.

## Author contributions

Authorship contributions are described based on the standard CRediT, Contributor Roles Taxonomy as follows”

Eva C Henningsen Data curation, Formal analysis, Investigation, Software, Visualization, Writing – original draft, Writing – review & editing

Eric S Nazareno Data curation, Investigation, Writing – review & editing

Rebecca Spanner Data curation, Investigation, Project administration, Writing – review & editing

David Lewis Investigation, Writing – review & editing Jibril Lubega Investigation, Writing – review & editing

Erin P Moreau Investigation, Writing – review & editing Brendan Boesen Supervision, Writing – review & editing

Eric Stone Funding acquisition, Supervision, Writing – review & editing Matt Moscou Funding acquisition, Supervision, Writing – review & editing

Kostya Kanyuka Conceptualization, Funding acquisition, Project administration, Resources, Supervision, Writing – review & editing

Shahryar Kianian Conceptualization, Funding acquisition, Resources, Writing – review & editing

Brian J Steffenson Conceptualization, Funding acquisition, Project administration, Resources, Supervision, Writing – review & editing

Peter N Dodds Conceptualization, Funding acquisition, Methodology, Supervision, Writing – original draft, Writing – review & editing

Jana Sperschneider Conceptualization, Formal analysis, Methodology, Software, Supervision, Writing – original draft, Writing – review & editing

Melania Figueroa Conceptualization, Funding acquisition, Methodology, Project administration, Supervision, Writing – original draft, Writing – review & editing

## Supplementary Material

**Supplementary Table 1.** Metadata for phased chromosome level assemblies of *Puccinia coronata* f. sp. *avenae*.

**Supplementary Table 2.** Virulence (H) or avirulence (L) of *Puccinia coronata* f. sp. *avenae* isolates with haplotype-phased genome references on a subset of 31 oat lines from the USA differential set.

**Supplementary Table 3.** Chromatin contacts between chromosomes for haplotypes from ten *Puccinia coronata* f. sp. *avenae* isolates.

**Supplementary Table 4.** Assembly and annotation statistics for the genome references of 10 *Puccinia coronata* f. sp. *avenae* isolates.

**Supplementary Table 5.** Repeat family coverage of chromosome scaffolds in hap45 and hap46 from *Puccinia coronata* f. sp. *avenae* isolate 94ISR005.

**Supplementary Table 6.** Mapping coverage and percent scaffold coverage for main scaffolds of 94ISR005.

**Supplementary Table 7.** Non-reference variant counts (including missing calls) derived from pangenome graph alignment of 52 *Puccinia coronata* f. sp. *avenae* haplotypes. Haplotypes in rows were used as the vcf reference, haplotypes in columns were the samples.

**Supplementary Table 8.** Percent coverage of recombination blocks of 52 *Puccinia coronata* f. sp. *avenae* haplotypes. Table is read as coverage of row haplotypes by sequences shared with column haplotypes. Colors reflect the range of coverage, with red being highest (96.79%) and white being lowest (0.00%).

**Supplementary Table 9.** *k*-mer containment results for 352 isolates of *Puccinia coronata* f. sp. *avenae* on 20 haplotypes generated in this study. Colors reflect range of values for percent identity and percent shared *k*-mers scaled separately. Red is highest (100% for both identity and shared *k*-mers) and white is lowest (99.01% for identity, 82.53% for shared *k*-mers).

**Supplementary Table 10.** *k*-mer containment for three described (Henningsen et al. 2024) and eight novel *b* locus alleles in short-read data for 352 Puccinia coronata f. sp. avenae isolates. Colors reflect the range of values for percent identity and percent shared *k*-mers scaled separately, with red being highest (100% for identity and shared *k*-mers) and white being lowest (90.77% for identity and 13.10% for shared *k*-mers).

**Supplementary Table 11.** Information about orthogroups derived from proteomes for 40 *Puccinia coronata* f. sp. *avenae* haplotypes, including the type (core versus non-core), number of alleles, number of secreted and effector members and expression of orthogroup members from isolate 21ACT116. CMC = core multiple-copy; CSC = core single-copy; private = only found in one haplotype.

**Supplementary Table 12.** Differential expression (DE), allele-specific expression (ASE), and allele-specific regulation (ASR) summaries for 21ACT116. The number of genes used for DE analysis was 38,618; the number of heterozygous loci used for ASE and ASR analysis was 9,419 (18,838 genes).

**Supplementary Fig. 1**. Heatmap of virulence for 25 *P. coronata* f. sp. *avenae* (*Pca*) isolates across 31 oat differential lines. Superscripts in isolate names indicate the source of previously published phenotype data (1 –Hewitt et al. 2024; 2 – Miller et al. 2020; 3 – Henningsen et al. 2022). Colors indicate virulence (red) and avirulence (yellow).

**Supplementary Fig. 2** Violin plots of the lengths (x-axis) for each of the 18 homologous chromosomes (y-axis) across 52 *Pca* haplotypes. Points represent chromosome lengths for individual haplotypes. Scaff1 is not shown as it is only found in hap46. The violin plot outline represents density across the distribution.

**Supplementary Fig. 3.** Synteny of *Pca* chromosomes (numbered 1-18, columns) from the 40 unique haplotypes (rows; excluding hap21, hap22, hap24, and haplotypes 41-44, haplotypes 47-50, hap52). Numbers on the left side reflect haplotype name (i.e. hap1 = 1, hap2 = 2). Order of haplotypes corresponds to their phylogenetic relationships. Colored lines between haplotypes correspond to the relationships between their sequences, including synteny (grey), inversion (gold), translocation (red) and duplication (blue).

**Supplementary Fig. 4.** Dotplot of genome alignment between chromosomes from hap35 (x-axis) and hap46 (y-axis).

**Supplementary Fig. 5.** Chromatin contact information supporting the phasing of Scaff1 into hap46. **A)** Heatmap of chromatin contact frequency along chromosomes from hap45 and hap46 (including Scaff1). Colors correspond to the range of contact frequency from high (red) to low (blue). **B)** Heatmap of chromatin contact frequency along Scaff1 from hap46. Colors correspond to the range of contact frequency from high (red) to low (blue). **C)** *trans* contact frequency of the main 18 chromosomes and extra scaffold (S1 = Scaff1) from *P. coronata* f. sp. *avenae* (*Pca*) 94ISR005 (hap45, hap46).

**Supplementary Fig. 6.** HiFi read coverage of three clonal *Pca* isolates (94ISR005, 94ISR022, 94ISR045) when mapped to 94ISR005 chromosomes (hap45, hap46). Coverage of 1 Kb bins for mappings including secondary (red) and primary only (black) are shown for chromosome 1 (chr1) from both haplotypes of 94ISR005 compared to Scaff1 which is only found in hap46.

**Supplementary Fig 7.** Heatmap of coverage for 2,384 repeat families (rows) across the 37 chromosomes (columns) from *P. coronata* f. sp. *avenae* isolate 94ISR005. Chromosomes from hap45 and hap46 are indicated with grey and white-filled boxes, respectively. Fill color indicates repeat family coverage, with white indicating no coverage and red indicating ≥ 1% coverage. S1 = Scaff1. Repeat families and chromosomes were clustered, with the chromosome clustering dendrogram shown at the top.

**Supplementary Fig 8.** Density plot of Kimura distances for LTR repetitive elements on chromosomes from hap45 and hap46 from *P. coronata* f. sp. *avenae* isolate 94ISR005. The density curve for Scaff1 is represented by a red line.

**Supplementary Fig. 9. A)** Recombination overlaps for *Pca* haplotype subgroups which share recombination blocks. The percentage of hap13, hap14, and hap51 which is covered by recombination blocks from the US population (left) or other population subgroups (middle, right) is shown at the bottom (% covered). **B)** Hypothesized introduction of hap14 (≈hap52) into the USA. Both haplotypes from CRO471 (hap51, hap52) are related to others from the global *Pca* population.

**Supplementary Fig. 10.** Tree of *STE3.2* alleles from Pca203 (hap1, hap2) and 20 additional *P. coronata* f. sp. *avenae* haplotypes rooted on the *Pgt STE3.2.1* allele. The tree scale is mean substitutions per site. Branches with bootstrap support ≥ 80% (100 cycles) are shown with white circles at the branch midpoint.

**Supplementary Fig. 11.** Protein trees for novel and characterized *b* locus alleles and schematic of variable, DNA binding, and conserved domains. Maximum Likelihood trees of **A)** *bW* and **B)** *bE* alleles identified in *P. coronata* f. sp. *avenae* (*Pca*) previously (Henningsen et al. 2024) with alleles from 20 new *Pca* haplotypes, rooted on the *Pgt bW1-HD1* and *Pgt bE1-HD2* alleles, respectively. Bootstraps from 100 cycles 80% and higher are shown with white circles at branch midpoints. Tree scales indicate mean substitutions per site. **C)** Diagram of bW and bE proteins from *Pca*, including variable, DNA binding (pink box) and conserved regions.

**Supplementary Fig. 12.** Pangenome openness assessed with cluster (orthogroups + singleton proteins) presence-absence variation for the proteomes from 40 unique haplotypes (excluding hap5, hap21, hap22, and haplotypes 41-44, haplotypes 47-50, hap52). Red curve is a Power Law fit. The value for the Heaps’ decay parameter α is shown, with α > 1 indicating a closed pangenome. The horizontal black dashed line is the pangenome size estimate derived from estimating the Chao lower bound (y = 36,053).

**Supplementary Fig. 13.** Summary of cluster membership and categories. **A)** Histogram of haplotypes per cluster constructed from the proteomes of 40 unique *Pca* haplotypes. **B)** Core and non-core proteome compartments, visualized as both a count of clusters (left) and a count of the proteins within the clusters (right). Fill reflects cluster classifications within the broader core and non-core categories. Abbreviations: CMC = core multiple copy; CSC = core single copy. **C)** Histogram of haplotypes represented per cluster for different categories of orthogroups (OGs). Top to bottom: non-secreted, secreted non-effector, effector. **D)** Percentage (y-axis) of OGs in different categories (columns, x-axis) and whether they are also categorized as non-secreted, secreted non-effector, or effector OGs (fill color). The total number of genes represented is shown at the top of each bar.

**Supplementary Fig. 14.** Differential expression visualized for 18,151 genes as log_2_(F old change) (x-axis) and -log(adjusted *p*-value) (y-axis) values across different orthogroup categories (rows and columns) when comparing **A)** infected tissue at four days post-inoculation (4dpi) with germinating spores (GS), **B)** infected tissue at six days post-inoculation (6dpi) with GS, and **C)** 6dpi versus 4dpi. Points represent genes, with point color reflecting whether they were upregulated (red), downregulated (blue), or not differentially expressed (grey). Thresholds for calling differential expression are shown as dashed lines (adjusted *p*-value < 0.05; fold change ≥ 2 or ≤ -2).

**Supplementary Fig. 15.** Percentage (y-axis) of genes from orthogroups in different categories (columns, x-axis) and whether they are expressed or not (top row) or for expressed genes, whether they were differentially regulated (bottom row) for the comparison of 4 days post-inoculation (4dpi) leaf samples to germinated spores (GS). Fill color corresponds to expression or regulation of genes: expressed (light grey), not expressed (black), downregulated (blue), not DE (grey), or upregulated (red). The total number of genes represented is shown at the top of each bar.

**Supplementary Fig. 16. A)** Percentage of heterozygous loci (y-axis) from 21ACT116 in each orthogroup category (x-axis, columns) exhibiting degrees of differential expression (fill color) in germinated spores (GS) and at 4 days post-inoculation (4dpi). Categories represent either expression/no expression or ranges for the absolute value of log_2_(Fold change): Not differentially expressed (DE) -0 to < 1), 1 to < 2); 2 to < 6); ≥ 6. Panels showing degree of DE include only expressed loci. **B)** Percentage of allele pairs that showed expression (top row) and percentage of expressed allele pairs exhibited allele-specific regulation (ASR) (bottom row) for different orthogroup categories (x-axis, columns) for the comparison of 4dpi infected leaf samples to GS. Fill color reflects what type of difference was observed (Same = both alleles had the same regulation pattern; One DE = one allele was DE, the other was not; Opposite = one allele was upregulated and the other downregulated). For panels **A** and **B**, the total number of genes represented is shown at the top of each bar.

