## Supplementary Figures for "Haplotype-resolved genomes of diverse oat crown rust isolates reveal both global dispersion of long-lived clonal haplotypes and limited recombination between haplotypes"

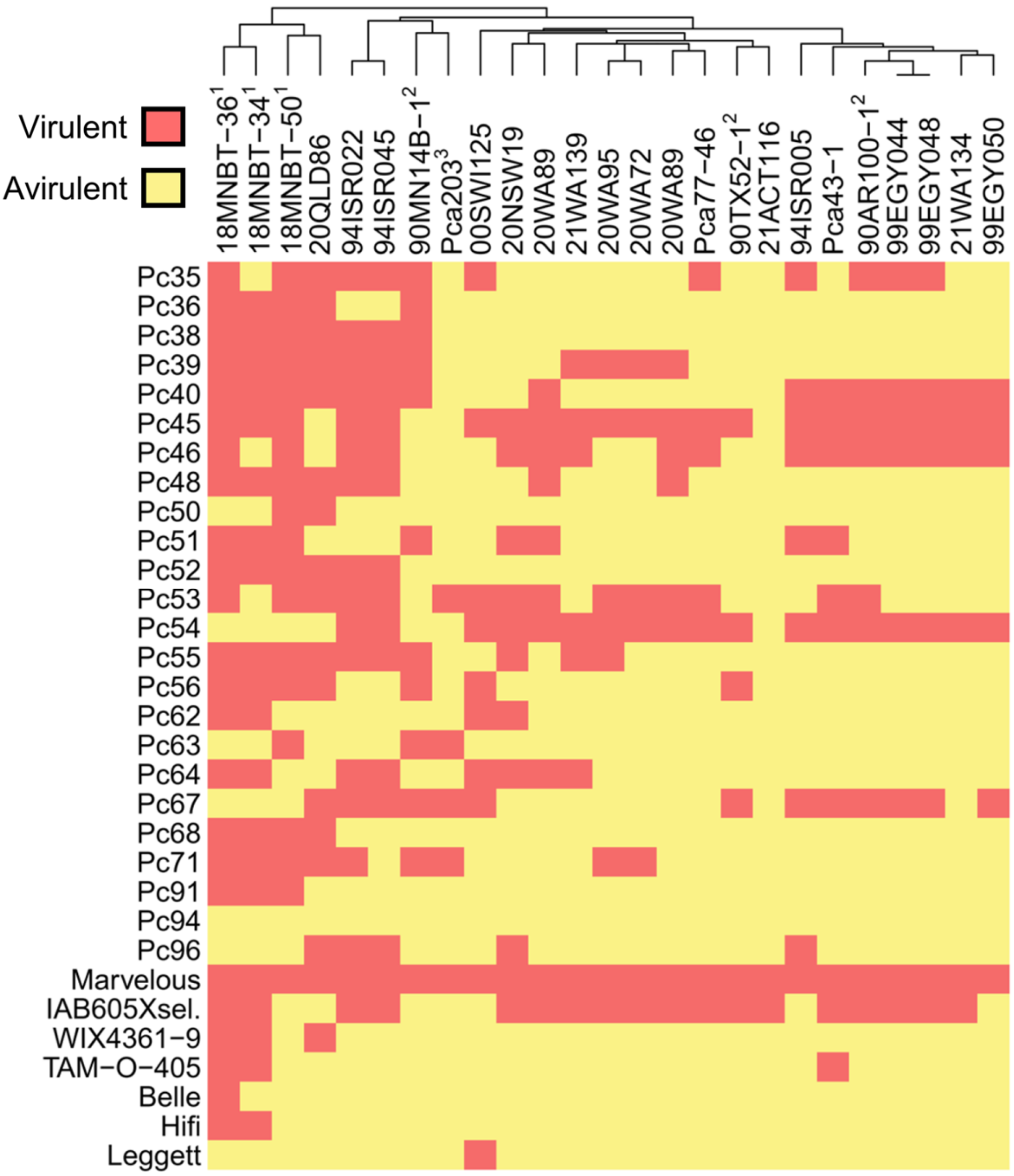


**Supplementary Fig. 1**. Heatmap of virulence for 25 *P. coronata* f. sp. *avenae* (*Pca*) isolates across 31 oat differential lines. Superscripts in isolate names indicate the source of previously published phenotype data (1 –Hewitt et al. 2024; 2 – Miller et al. 2020; 3 – Henningsen et al. 2022). Colors indicate virulence (red) and avirulence (yellow).

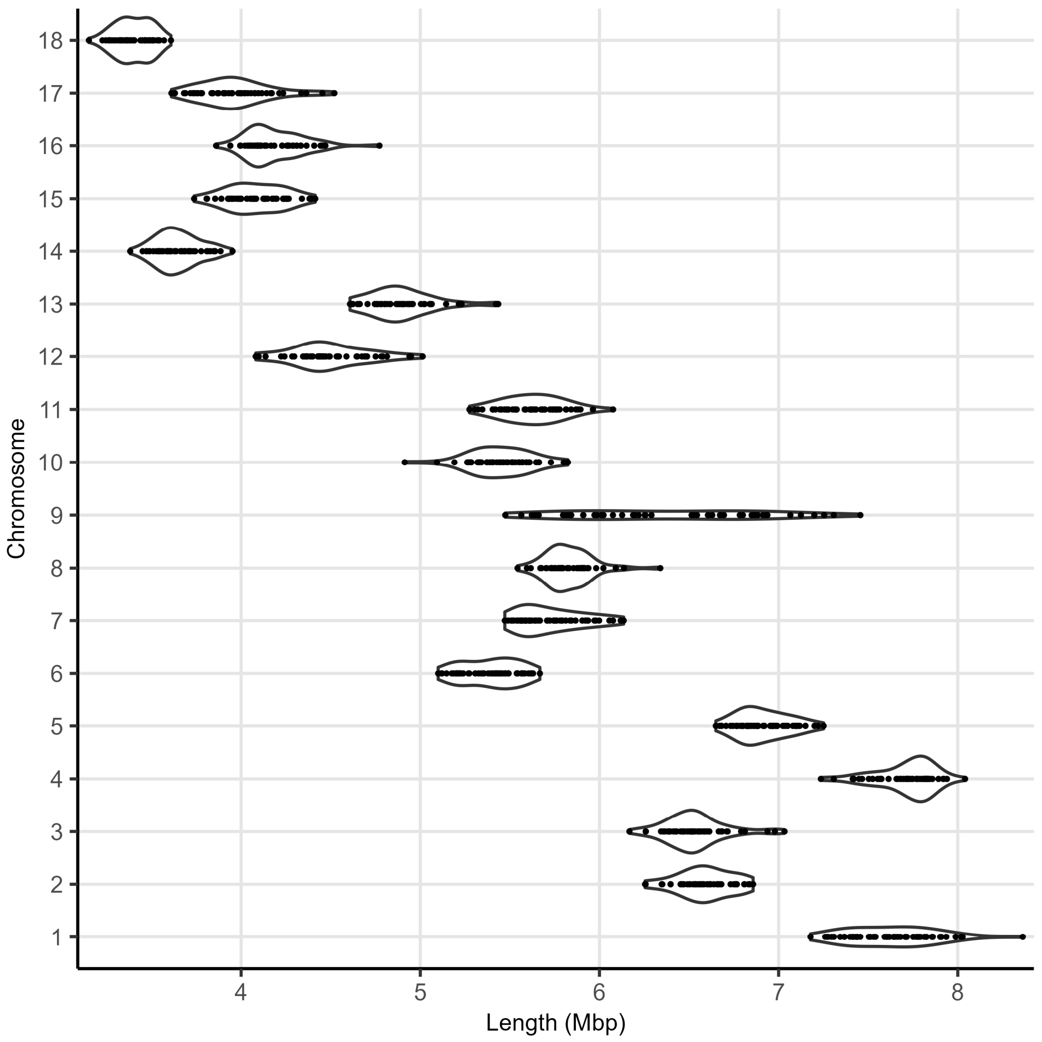


**Supplementary Fig. 2** Violin plots of the lengths (x-axis) for each of the 18 homologous chromosomes (y-axis) across 52 *Pca* haplotypes. Points represent chromosome lengths for individual haplotypes. Scaff1 is not shown as it is only found in hap46. The violin plot outline represents density across the distribution.

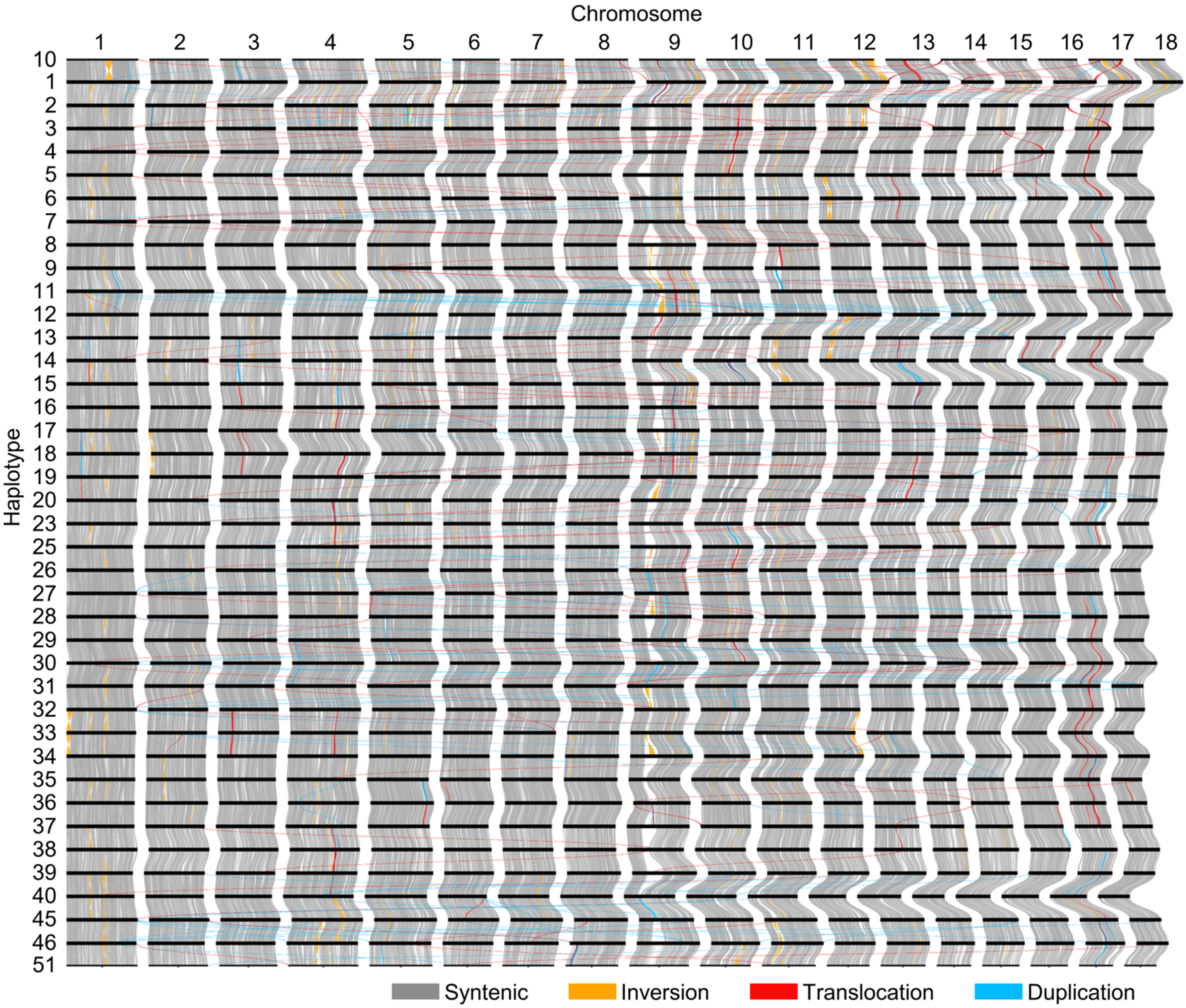


**Supplementary Fig. 3.** Synteny of *Pca* chromosomes (numbered 1-18, columns) from the 40 unique haplotypes (rows; excluding hap21, hap22, hap24, and haplotypes 41-44, haplotypes 47-50, hap52). Numbers on the left side reflect haplotype name (i.e. hap1 = 1, hap2 = 2). Order of haplotypes corresponds to their phylogenetic relationships. Colored lines between haplotypes correspond to the relationships between their sequences, including synteny (grey), inversion (gold), translocation (red) and duplication (blue).

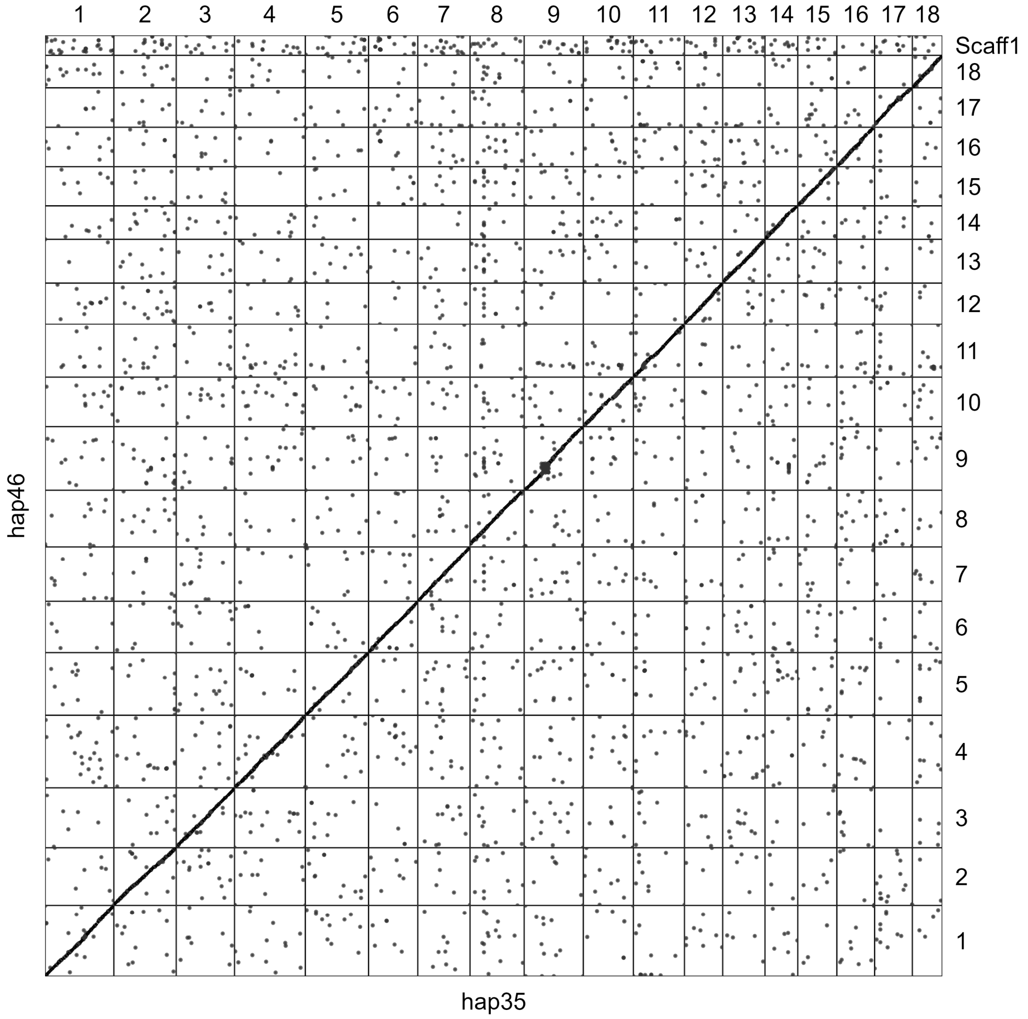


**Supplementary Fig. 4.** Dotplot of genome alignment between chromosomes from hap35 (x-axis) and hap46 (y-axis).

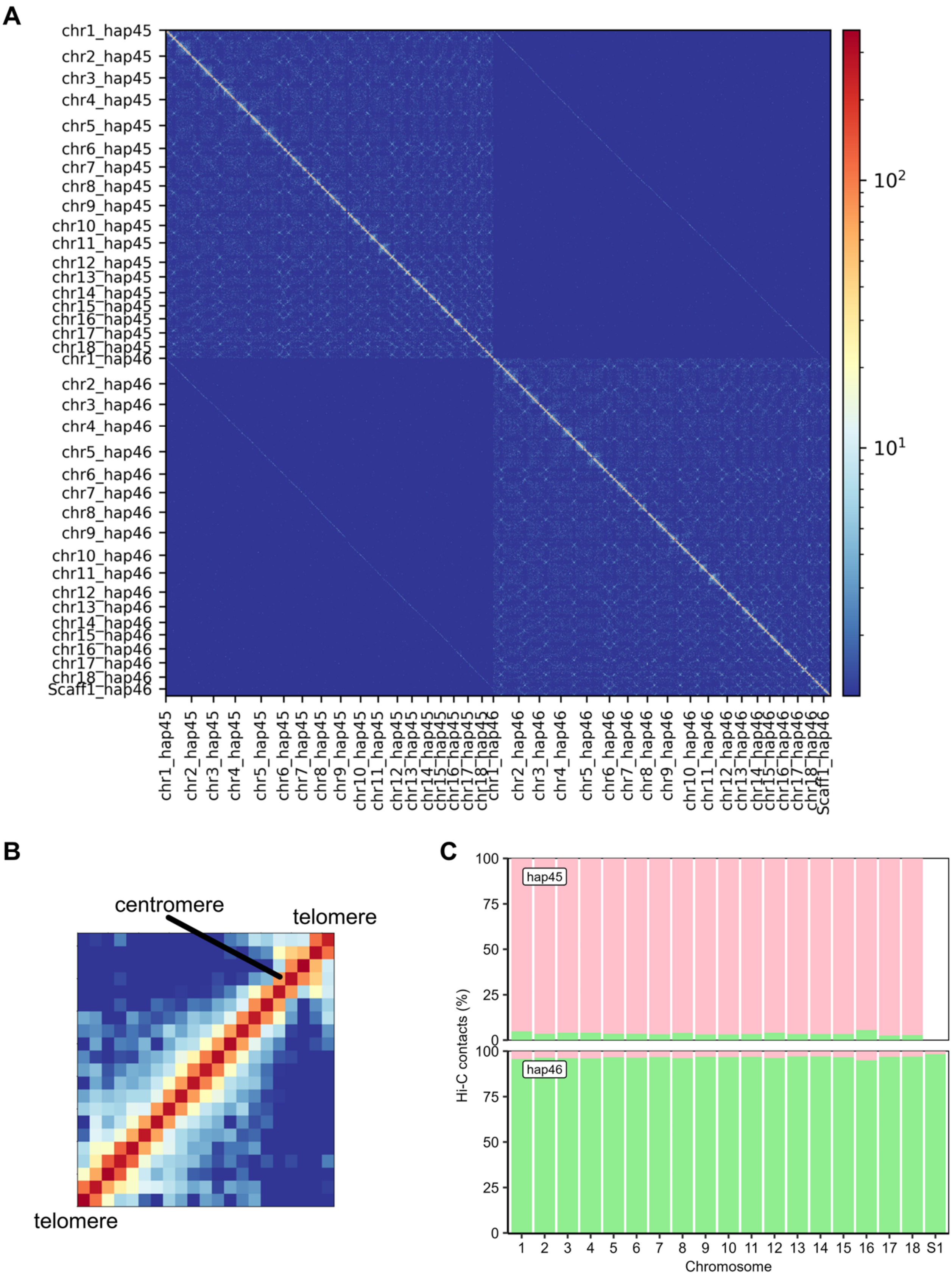


**Supplementary Fig. 5.** Chromatin contact information supporting the phasing of Scaff1 into hap46. **A)** Heatmap of chromatin contact frequency along chromosomes from hap45 and hap46 (including Scaff1). Colors correspond to the range of contact frequency from high (red) to low (blue). **B)** Heatmap of chromatin contact frequency along Scaff1 from hap46. Colors correspond to the range of contact frequency from high (red) to low (blue). **C)** *trans* contact frequency of the main 18 chromosomes and extra scaffold (S1 = Scaff1) from *P. coronata* f. sp. *avenae* (*Pca*) 94ISR005 (hap45, hap46).

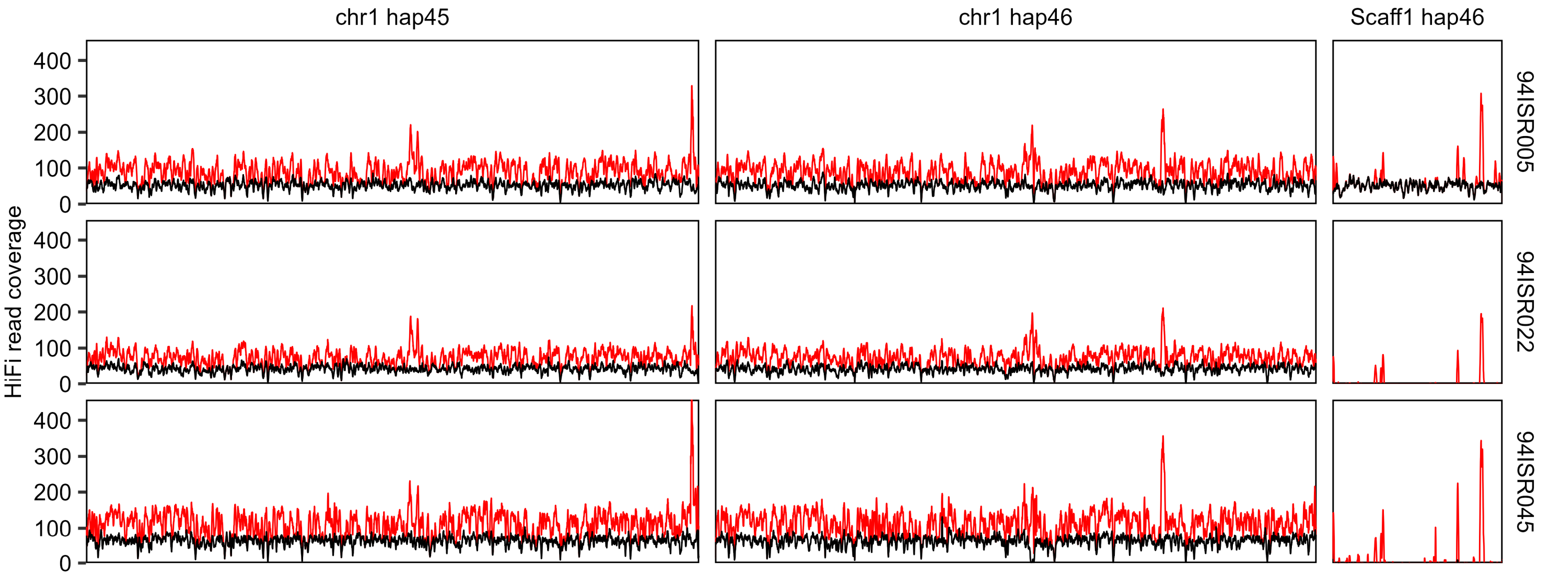


**Supplementary Fig. 6.** HiFi read coverage of three clonal *Pca* isolates (94ISR005, 94ISR022, 94ISR045) when mapped to 94ISR005 chromosomes (hap45, hap46). Coverage of 1 Kb bins for mappings including secondary (red) and primary only (black) are shown for chromosome 1 (chr1) from both haplotypes of 94ISR005 compared to Scaff1 which is only found in hap46.

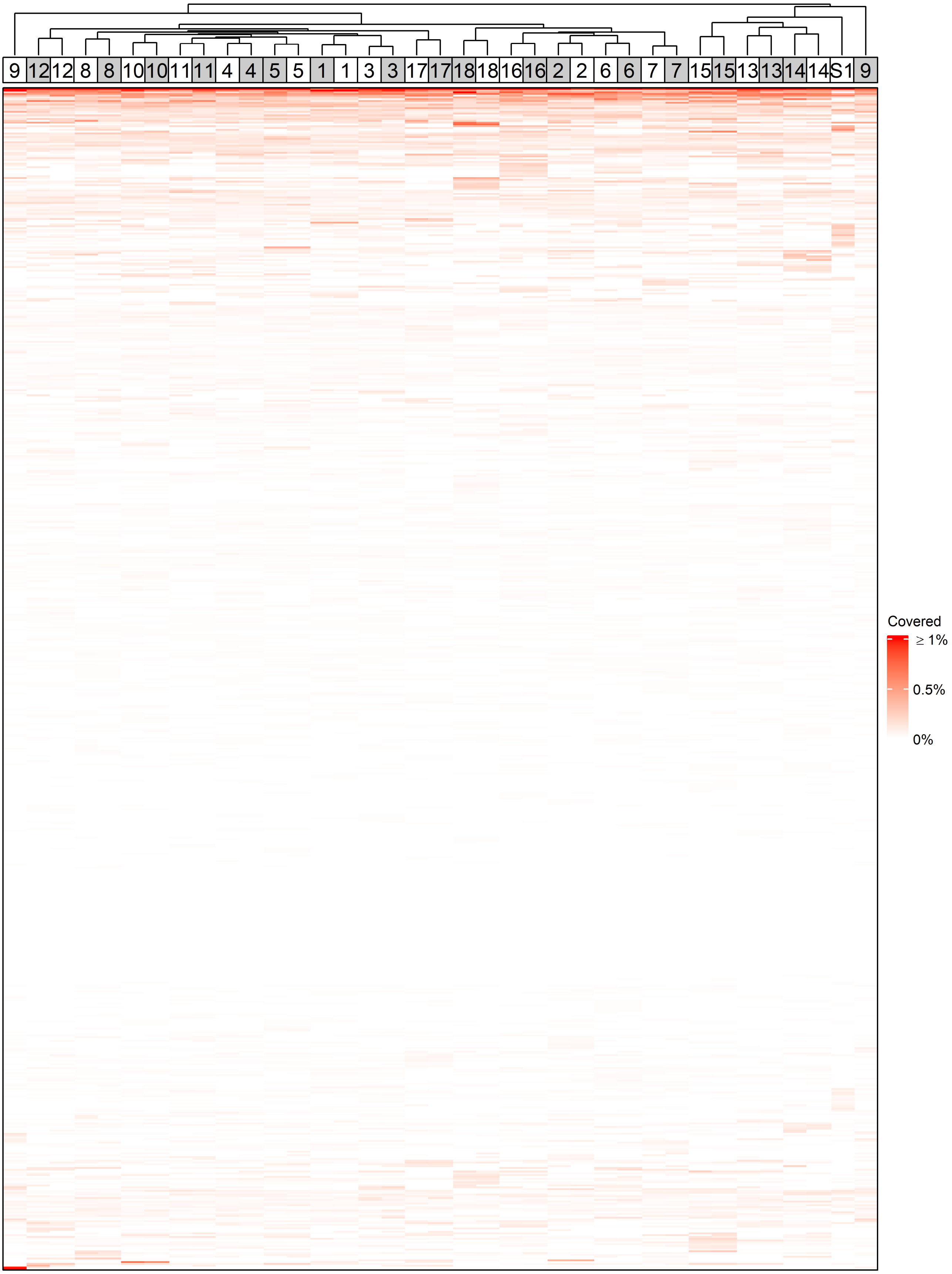


**Supplementary Fig 7.** Heatmap of coverage for 2,384 repeat families (rows) across the 37 chromosomes (columns) from *P. coronata* f. sp. *avenae* isolate 94ISR005. Chromosomes from hap45 and hap46 are indicated with grey and white-filled boxes, respectively. Fill color indicates repeat family coverage, with white indicating no coverage and red indicating ≥ 1% coverage. S1 = Scaff1. Repeat families and chromosomes were clustered, with the chromosome clustering dendrogram shown at the top.

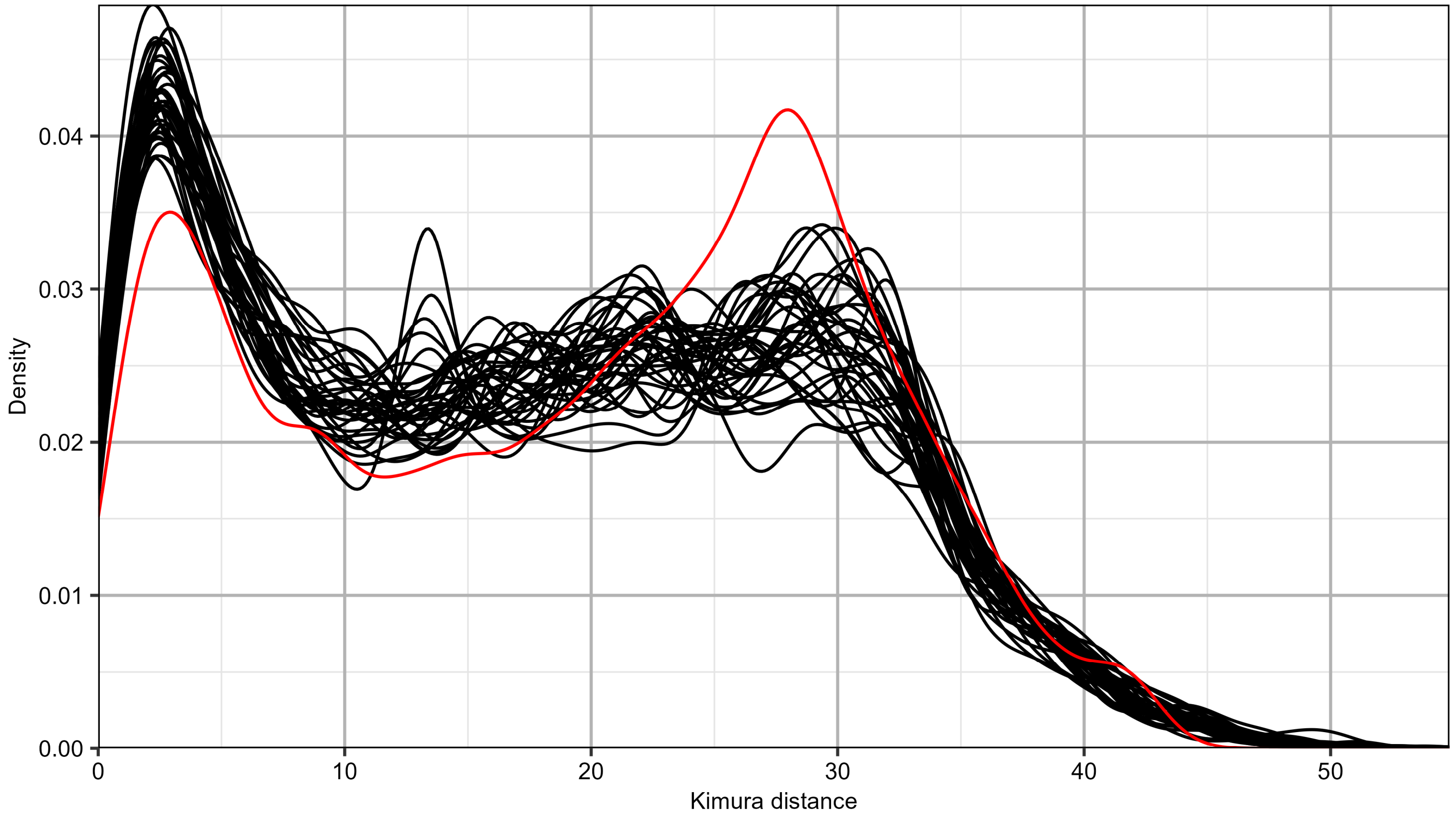


**Supplementary Fig 8.** Density plot of Kimura distances for LTR repetitive elements on chromosomes from hap45 and hap46 from *P. coronata* f. sp. *avenae* isolate 94ISR005. The density curve for Scaff1 is represented by a red line.

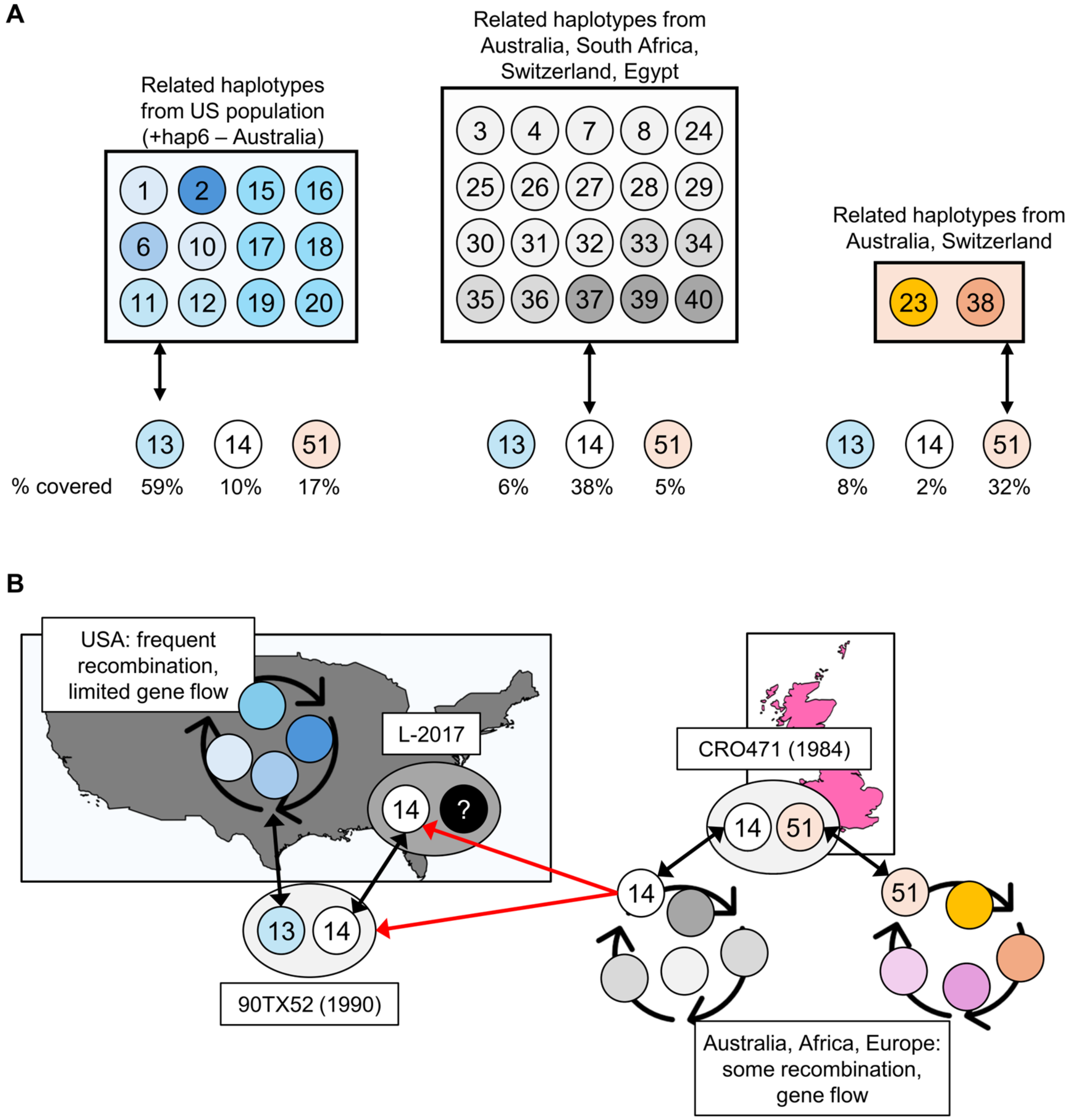


**Supplementary Fig. 9. A)** Recombination overlaps for *Pca* haplotype subgroups which share recombination blocks. The percentage of hap13, hap14, and hap51 which is covered by recombination blocks from the US population (left) or other population subgroups (middle, right) is shown at the bottom (% covered). **B)** Hypothesized introduction of hap14 (≈hap52) into the USA. Both haplotypes from CRO471 (hap51, hap52) are related to others from the global *Pca* population.

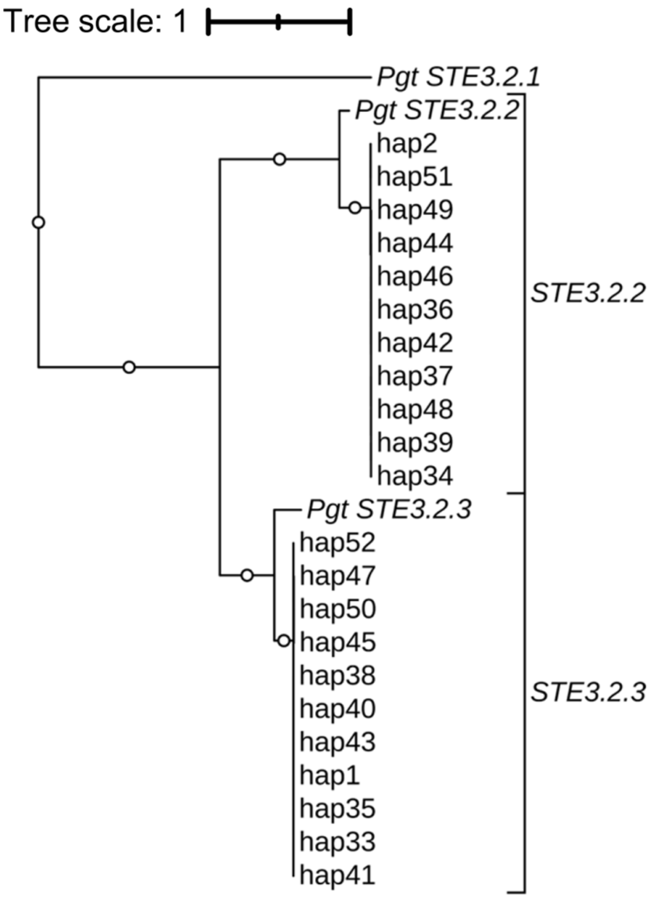


**Supplementary Fig. 10.** Tree of *STE3.2* alleles from Pca203 (hap1, hap2) and 20 additional *P. coronata* f. sp. *avenae* haplotypes rooted on the *Pgt STE3.2.1* allele. The tree scale is mean substitutions per site. Branches with bootstrap support ≥ 80% (100 cycles) are shown with white circles at the branch midpoint.

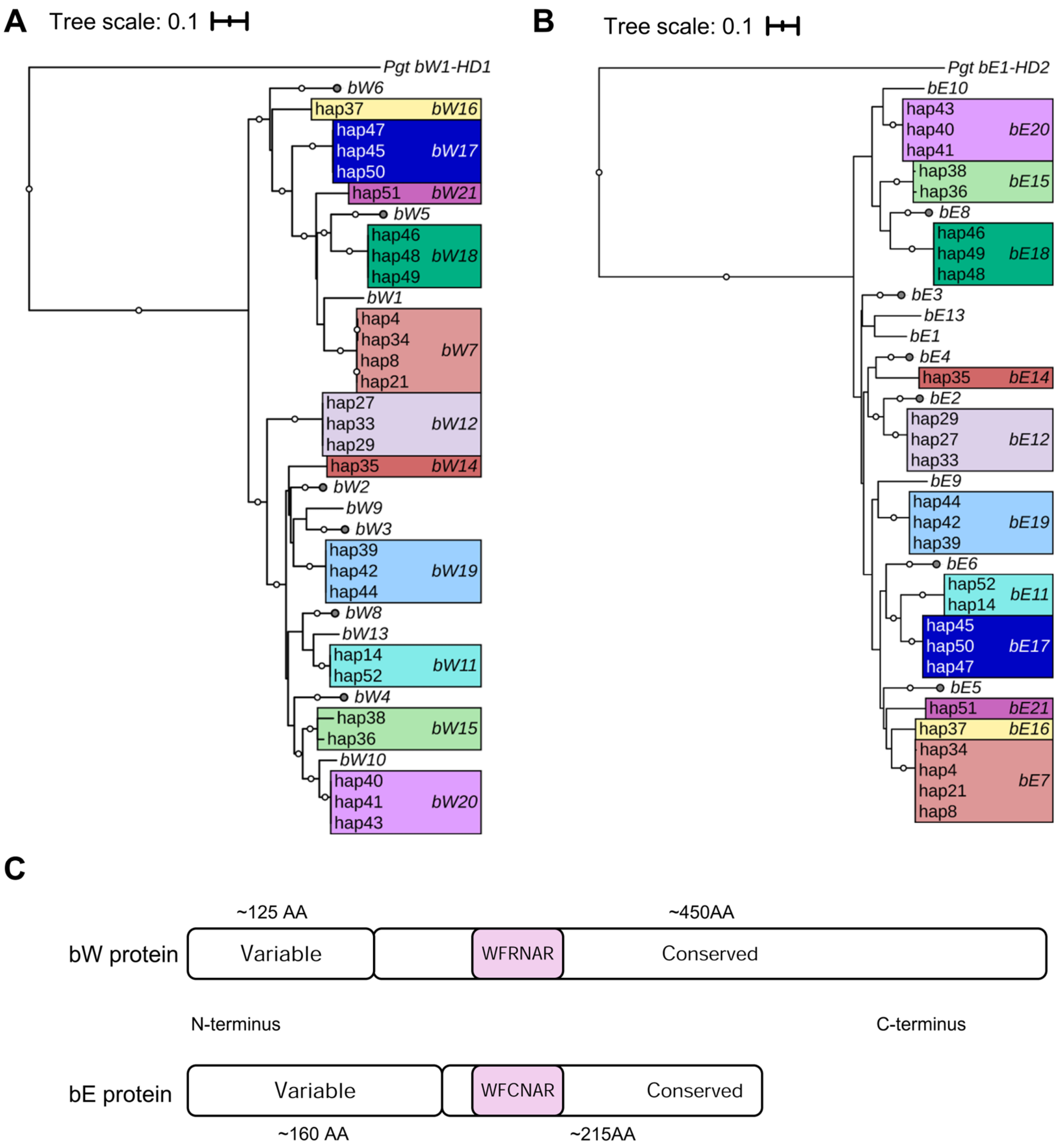


**Supplementary Fig. 11.** Protein trees for novel and characterized *b* locus alleles and schematic of variable, DNA binding, and conserved domains. Maximum Likelihood trees of **A)** *bW* and **B)** *bE* alleles identified in *P. coronata* f. sp. *avenae* (*Pca*) previously (Henningsen et al. 2024) with alleles from 20 new *Pca* haplotypes, rooted on the *Pgt bW1-HD1* and *Pgt bE1-HD2* alleles, respectively. Bootstraps from 100 cycles 80% and higher are shown with white circles at branch midpoints. Tree scales indicate mean substitutions per site. **C)** Diagram of bW and bE proteins from *Pca*, including variable, DNA binding (pink box) and conserved regions.

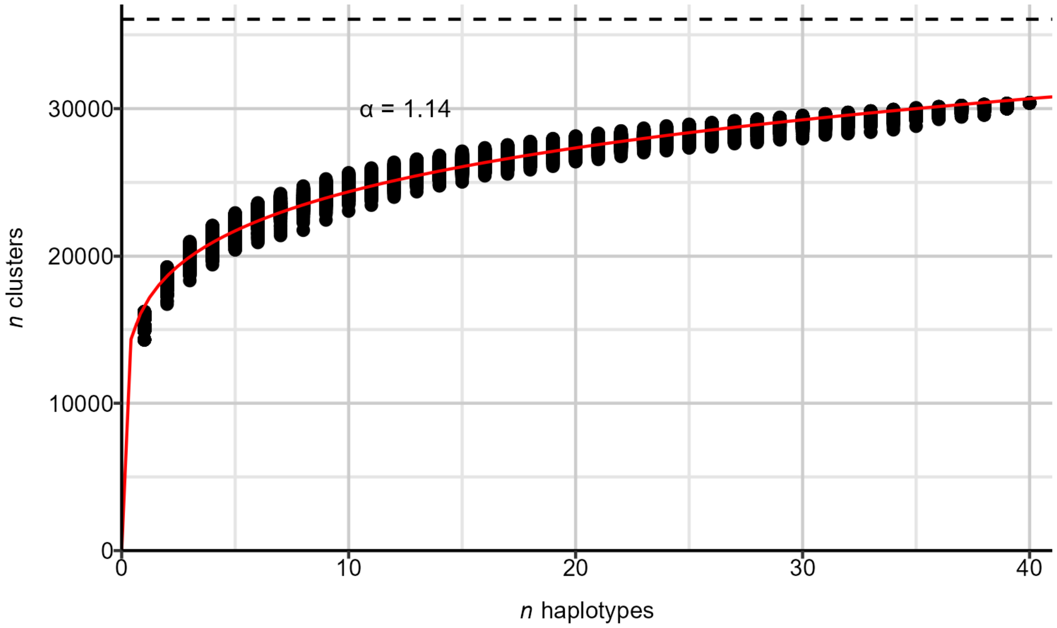


**Supplementary Fig. 12.** Pangenome openness assessed with cluster (orthogroups + singleton proteins) presence-absence variation for the proteomes from 40 unique haplotypes (excluding hap5, hap21, hap22, and haplotypes 41-44, haplotypes 47-50, hap52). Red curve is a Power Law fit. The value for the Heaps' decay parameter α is shown, with α > 1 indicating a closed pangenome. The horizontal black dashed line is the pangenome size estimate derived from estimating the Chao lower bound (y = 36,053).

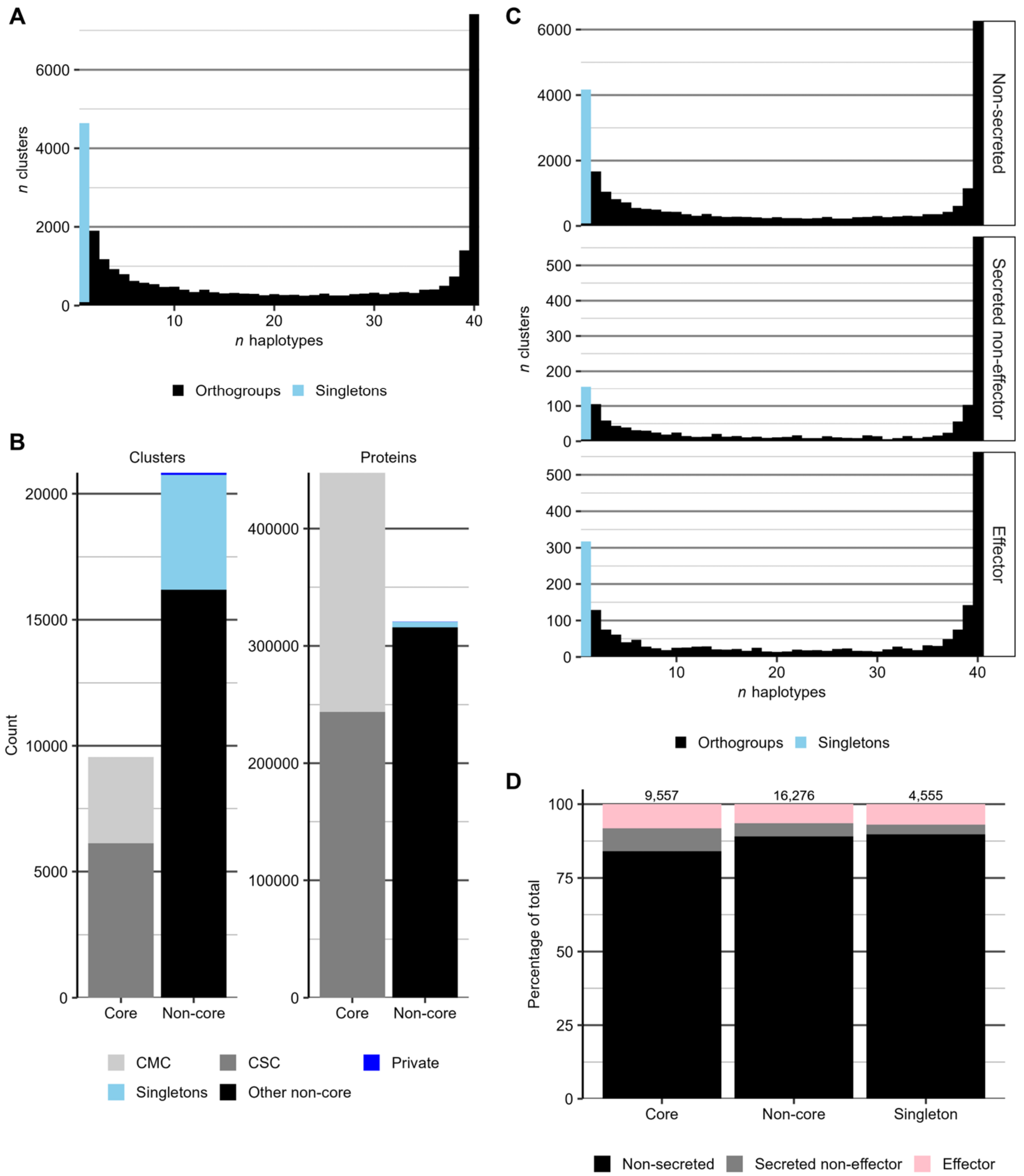


**Supplementary Fig. 13.** Summary of cluster membership and categories. **A)** Histogram of haplotypes per cluster constructed from the proteomes of 40 unique *Pca* haplotypes. **B)** Core and non-core proteome compartments, visualized as both a count of clusters (left) and a count of the proteins within the clusters (right). Fill reflects cluster classifications within the broader core and non-core categories. Abbreviations: CMC = core multiple copy; CSC = core single copy. **C)** Histogram of haplotypes represented per cluster for different categories of orthogroups (OGs). Top to bottom: non-secreted, secreted non-effector, effector. **D)** Percentage (y-axis) of OGs in different categories (columns, x-axis) and whether they are also categorized as non-secreted, secreted non-effector, or effector OGs (fill color). The total number of genes represented is shown at the top of each bar.

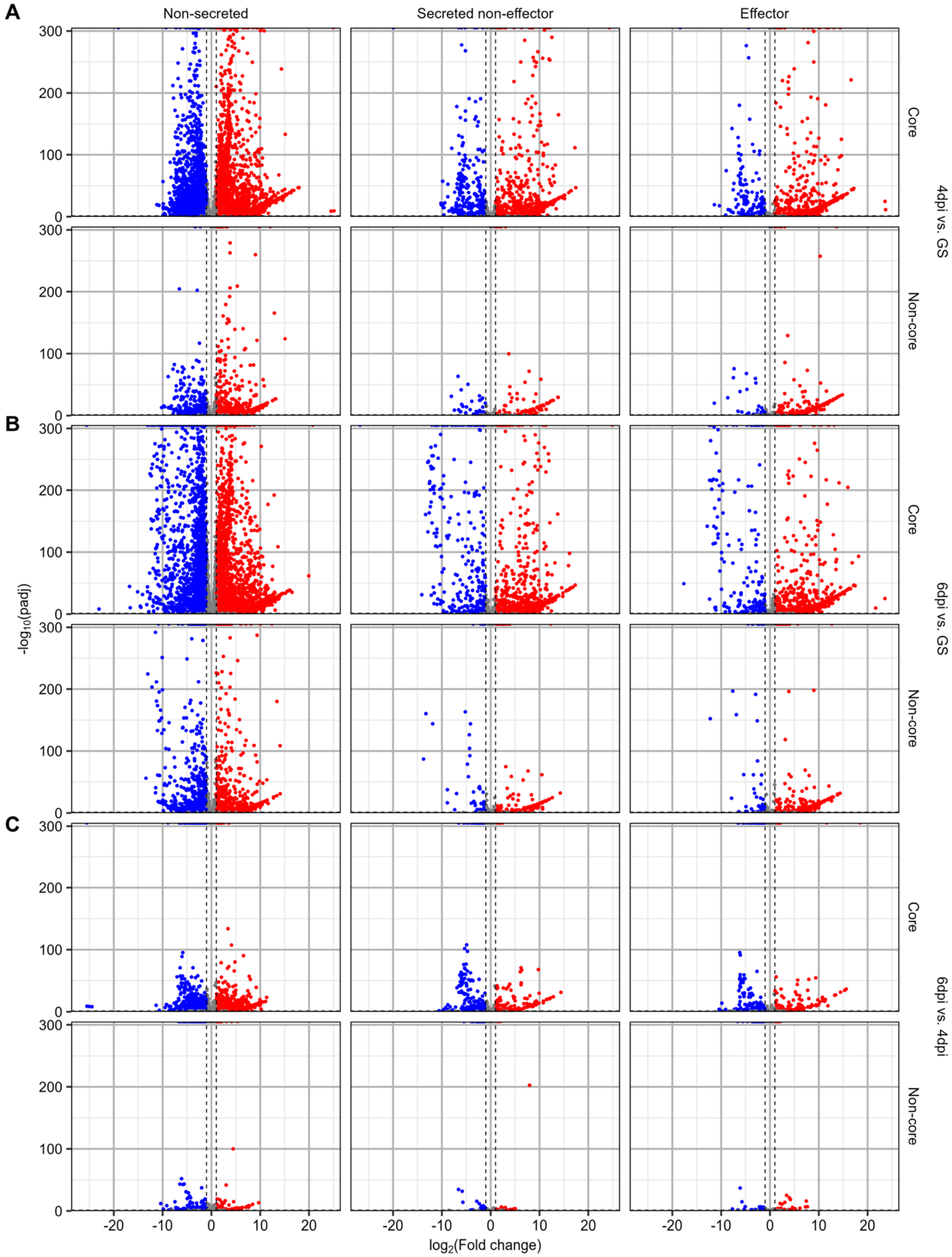


**Supplementary Fig. 14.** Differential expression visualized for 18,151 genes as $\log_2$(F old change) (x-axis) and -log(adjusted *p*-value) (y-axis) values across different orthogroup categories (rows and columns) when comparing **A)** infected tissue at four days post-inoculation (4dpi) with germinating spores (GS), **B)** infected tissue at six days post-inoculation (6dpi) with GS, and **C)** 6dpi versus 4dpi. Points represent genes, with point color reflecting whether they were upregulated (red), downregulated (blue), or not differentially expressed (grey). Thresholds for calling differential expression are shown as dashed lines (adjusted *p*-value < 0.05; fold change ≥ 2 or ≤ -2).

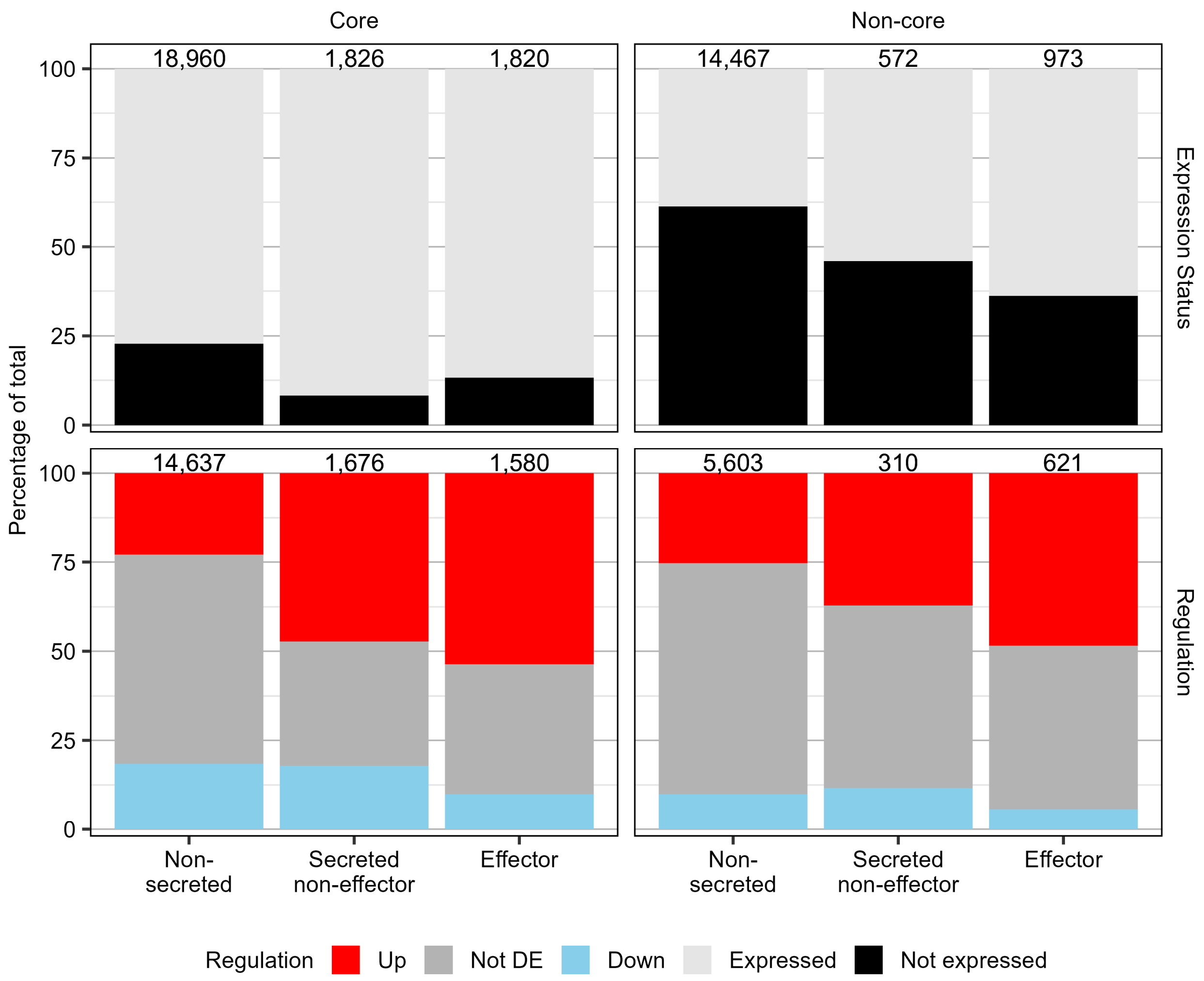


**Supplementary Fig. 15.** Percentage (y-axis) of genes from orthogroups in different categories (columns, x-axis) and whether they are expressed or not (top row) or for expressed genes, whether they were differentially regulated (bottom row) for the comparison of 4 days post-inoculation (4dpi) leaf samples to germinated spores (GS). Fill color corresponds to expression or regulation of genes: expressed (light grey), not expressed (black), downregulated (blue), not DE (grey), or upregulated (red). The total number of genes represented is shown at the top of each bar.

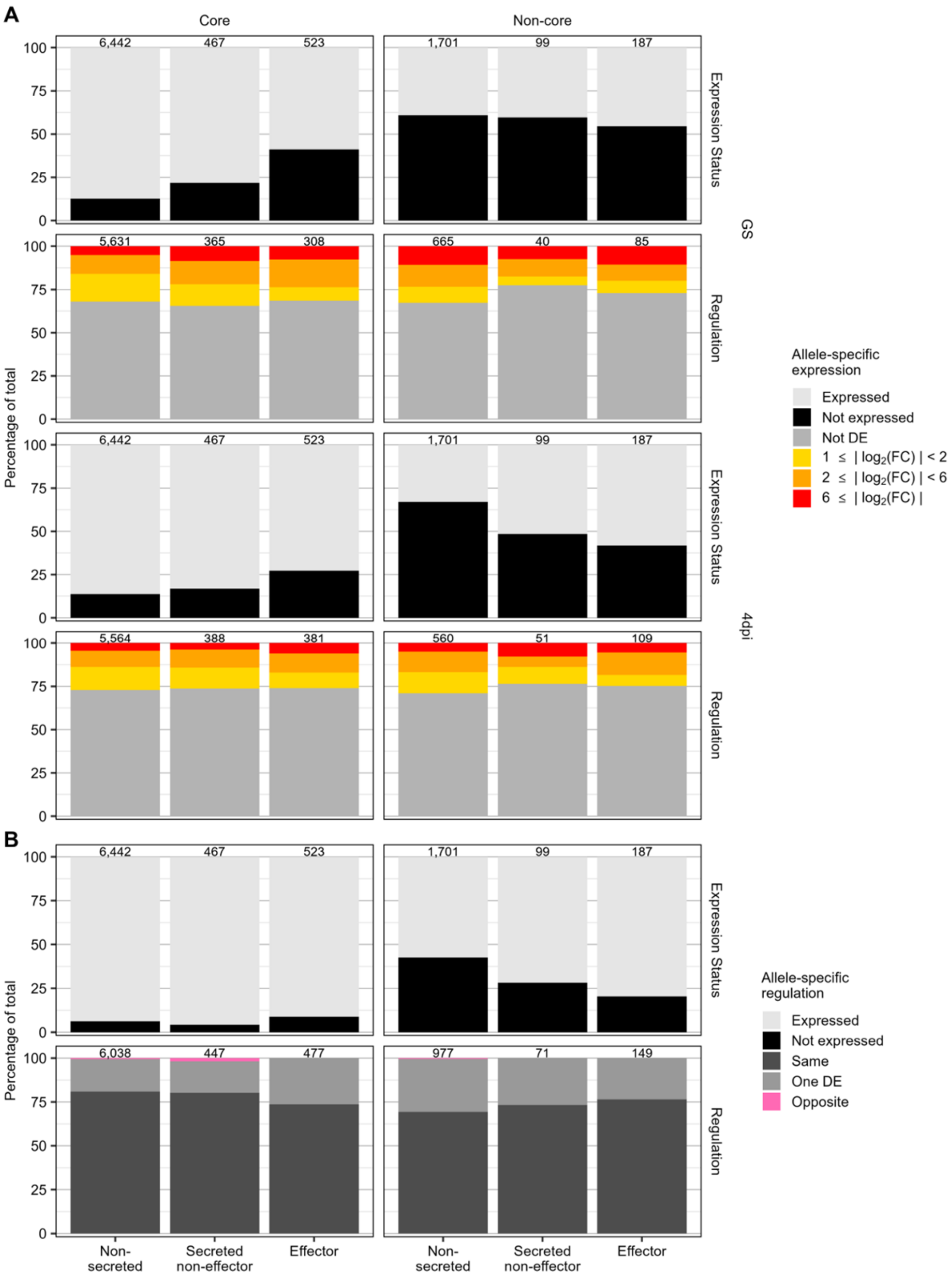


**Supplementary Fig. 16. A)** Percentage of heterozygous loci (y-axis) from 21ACT116 in each orthogroup category (x-axis, columns) exhibiting degrees of differential expression (fill color) in germinated spores (GS) and at 4 days post-inoculation (4dpi). Categories represent either expression/no expression or ranges for the absolute value of $log_2$(Fold change): Not differentially expressed (DE) - [0,1), [1,2); [2,6); ≥ 6. Panels showing degree of DE include only expressed loci. **B)** Percentage of allele pairs that showed expression (top row) and percentage of expressed allele pairs exhibited allele-specific regulation (ASR) (bottom row) for different orthogroup categories (x-axis, columns) for the comparison of 4dpi infected leaf samples to GS. Fill color reflects what type of difference was observed (Same = both alleles had the same regulation pattern; One DE = one allele was DE, the other was not; Opposite = one allele was upregulated and the other downregulated). For panels **A** and **B**, the total number of genes represented is shown at the top of each bar.
